# Prefrontal solution to the bias-variance tradeoff during reinforcement learning

**DOI:** 10.1101/2020.12.23.424258

**Authors:** Dongjae Kim, Jaeseung Jeong, Sang Wan Lee

**Affiliations:** Department of Bio and Brain Engineering, Korea Advanced Institute of Science and Technology (KAIST) Daejeon, 34141 Republic of Korea; Program of Brain and Cognitive engineering, Korea Advanced Institute of Science and Technology (KAIST) Daejeon, 34141 Republic of Korea; KAIST Center for Neuroscience-inspired AI, Korea Advanced Institute of Science and Technology (KAIST) Daejeon, 34141 Republic of Korea; KI for Health Science and Technology, Korea Advanced Institute of Science and Technology (KAIST) Daejeon, 34141 Republic of Korea; KI for Artificial Intelligence, Korea Advanced Institute of Science and Technology (KAIST) Daejeon, 34141 Republic of Korea

## Abstract

The goal of learning is to maximize future rewards by minimizing prediction errors. Evidence have shown that the brain achieves this by combining model-based and model-free learning. However, the prediction error minimization is challenged by a bias-variance tradeoff, which imposes constraints on each strategy’s performance. We provide new theoretical insight into how this tradeoff can be resolved through the adaptive control of model-based and model-free learning. The theory predicts the baseline correction for prediction error reduces the lower bound of the bias–variance error by factoring out irreducible noise. Using a Markov decision task with context changes, we showed behavioral evidence of adaptive control. Model-based behavioral analyses show that the prediction error baseline signals context changes to improve adaptability. Critically, the neural results support this view, demonstrating multiplexed representations of prediction error baseline within the ventrolateral and ventromedial prefrontal cortex, key brain regions known to guide model-based and model-free learning.

**One sentence summary:** A theoretical, behavioral, computational, and neural account of how the brain resolves the bias-variance tradeoff during reinforcement learning is described.

## Introduction

Reinforcement learning (RL) refers to the process of finding an optimal action policy to maximize cumulative rewards (Sutton and Barto, 1998). To develop this policy, the RL agent computes the prediction error, the discrepancy between an actual outcome from the environment and a predicted outcome. Abundant evidence has demonstrated the importance of prediction error signals in RL at the neural level (Cooper et al., 2012; Gläscher et al., 2012; Schultz, 1998).

Accumulating evidence has suggested that the brain uses two distinctly different strategies. One approach, called model-free (MF) RL, is to use a prediction error of the rewards, i.e., a reward prediction error (RPE), to find a policy that best suits the current situation. The other approach, called model-based (MB) RL, uses the prediction error of the state-transition, i.e., a state prediction error (SPE), that encodes information about the structure of the environment.

MF RL learns a policy without explicit knowledge about the task structure (Barto et al., 1983; Mnih et al., 2015, 2016; Watkins and Dayan, 1992). It is a computationally simple process, exhibiting a highly consistent behavior, i.e., a low variance. However, this does not outweigh a drawback in that its policy is often highly biased, and thus it may degenerate when facing a complex task structure or rapid changes in context (the leftmost side of Figure 1A). By contrast, MB RL has certain advantages over its counterpart, such as a faster adaptation speed (Kuvayev and Sutton, 1997) and behavioral flexibility. However, the optimistic view is dampened by the computational complexity and potential high bias error associated with it. Relying solely on the MB RL strategy may lead to a high bias or variance error in certain situations, for example, when the task structure is associated with an incorrect assumption or is too noisy to learn (Lengyel and Dayan, 2008) (the cases shown in the center of Figure 1A). The different performance constraints of MB and MF RL necessitate a combination of these two learning strategies.

**Figure 1.**
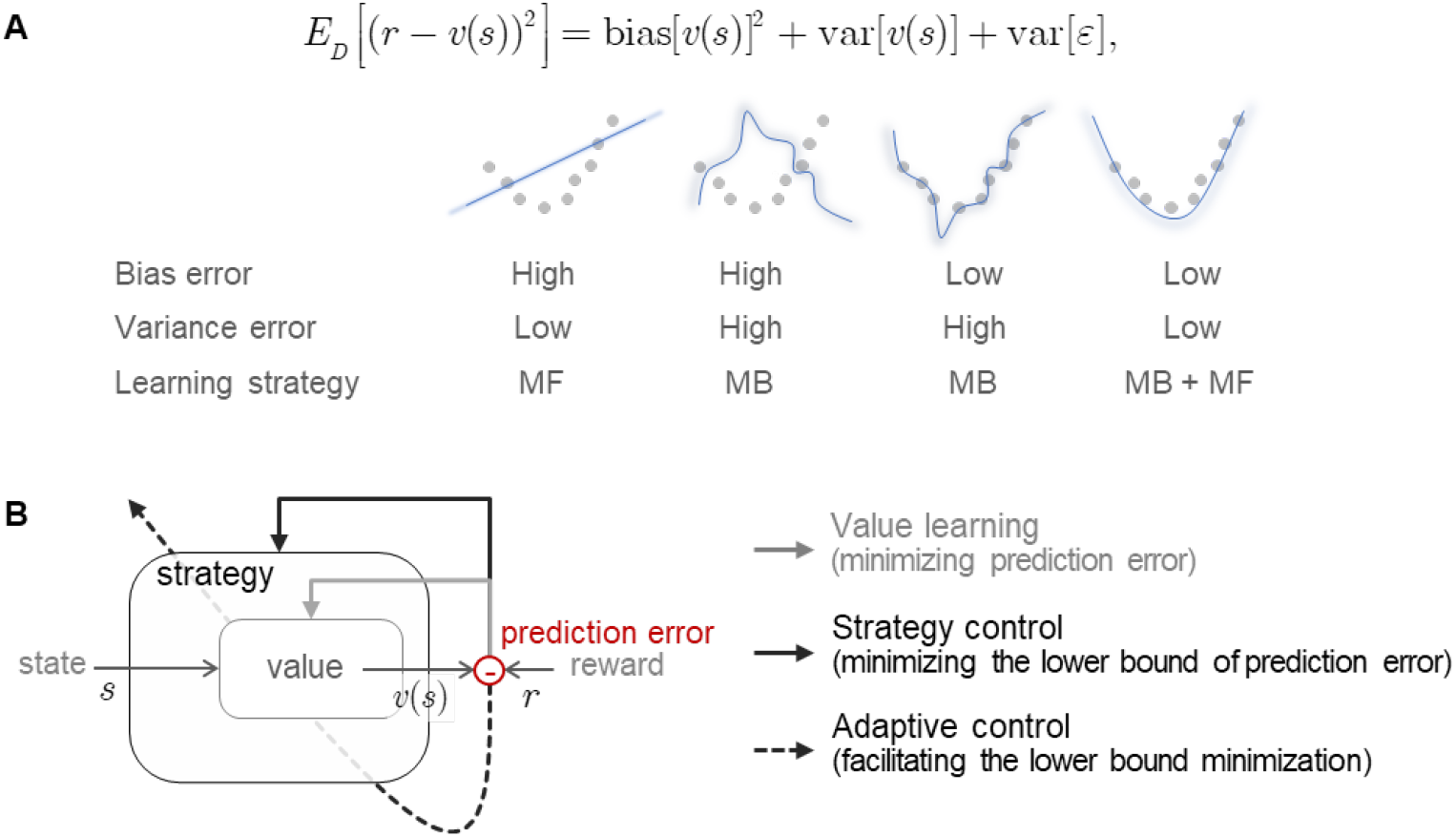
Model-based and model-free learning to resolve the bias-variance tradeoff. **(A)** The reinforcement learning problem is specified as finding a value function v(s) that minimizes the prediction error, and following the machine learning convention, this error can be decomposed into a bias error, variance error, and irreducible error arising from environmental noise. The choice of different learning strategies, such as model-free(MF) or model-based (MB) learning, can entail different types of bias-variance error (left three columns). It naturally follows that, for a given context, an ideal mixture of MB and MF exists that resolves the bias-variance error issue. **(B)** A conceptual learning framework for resolving the bias-variance tradeoff. Ideally, the agent minimizes the amount of prediction error (value learning) by combining MB and MF strategies (strategy control), leading to a reduction of both the bias and variance error (lower bound of the prediction error). The lower bound can be further reduced using a baseline correction for the prediction error (adaptive control).

There is ample evidence to support this idea (Balleine and Dickinson, 1998; Balleine and O’Doherty, 2010; Daw et al., 2005; Dolan and Dayan, 2013; Doya, 1999). Neuroimaging studies have also provided evidence of the mixed representation of MF and MB RL by identifying BOLD correlates of RPE (Cooper et al., 2012; Gläscher et al., 2012; Schultz, 1998) and SPE (Gläscher et al., 2010; Lee et al., 2014). Key variables for determining which strategy should guide the behavior at each moment in time are the prediction uncertainty (Daw et al., 2005) and reliability (Kim et al., 2019; Lee et al., 2014), which can be read from the RPE and SPE. This idea has been implemented using a threshold to classify the prediction error into a zero or non-zero prediction error, which dictate whether or not the “mistake” should be admitted. For example, repeated non-zero prediction error (e.g., RPE or SPE) casts doubt on the reliability of the current learning strategy (e.g., MF or MB), motivating the learning agent to explore alternative strategy.

Despite these attempts, little is known about how the combination of MB and MF RL helps reduce both the bias and variance error. This is a challenging task because minimizing the bias and variance error often conflict. Minimizing bias error can be achieved by increasing the capacity for adaptation, but it may increase behavioral sensitivity. On the other hand, minimizing variance error can be achieved by increasing behavioral sensitivity, but it comes at the cost of increasing risk of underfitting. Although the idea that the brain can solve the bias–variance tradeoff is not untenable, as recently discussed in a few contexts, such as Pavlovian-instrumental learning (Dorfman and Gershman, 2019), inference in an unpredictable environment (Glaze et al., 2018), and MB and MF learning (Filipowicz et al., 2020), it is difficult to quantify the degree of contribution of MB and MF to the bias–variance tradeoff, as indicated by both our earlier discussion (Figure 1A) and a recent study (Filipowicz et al., 2020).

The contributions of this study are as follows. First, we built a theoretical framework to show that even in the presence of environmental context changes, a tradeoff between bias and variance prediction error can be effectively solved through adaptive control of model-based and model-free learning, particularly when equipped with a baseline correction tracking the lower bound of the prediction error. Second, we used a Markov decision task with context changes allowing for this test and showing evidence of behavioral control contingent on the changes in context. This is further supported by our computational model with the baseline correction of the prediction error. Third, using this model, we examined the role of baseline correction in context-sensitive adaptive control. We showed that the baseline estimation reflects changes in context and the adaptation performance. Fourth, we showed the effect of baseline correction on the two areas in the prefrontal cortex, i.e., the lateral prefrontal cortex implementing arbitration control between model-based and model-free learning and the ventromedial prefrontal cortex implementing value learning. To further examine how these brain regions interact to effectively combine model-based and model-free learning, we ran functional connectivity analyses.

## Results

### New insight into bias-variance tradeoff in the context of model-based and mode-free learning

Here, we situate the bias-variance tradeoff, one of the most fundamental issues in model prediction, in the context of a dynamic environment. We decompose this into three problems, and for each we discuss a strategy within the framework of model-based and model-free learning.

### Value learning for bias-variance error minimization

Suppose that the reward function, *r* = *f*(*s*) + *ε*, where f is the true reward function and epsilon is zero-mean environment noise. The goal of reinforcement learning is to find a value function *v*(*s*) that represents the expected amount of future reward by minimizing the prediction error. The evaluation function for this minimization problem can be written as the average prediction error:

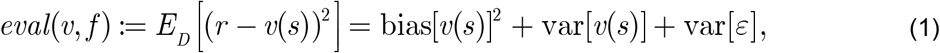

where bias[*v*(*s*)] = E_*D*_ [*v*(*s*)] − *f* (*s*) and var[*v*(*s*)] refers to the variance of *v*(*s*) (Figure 1A). Here, var[*ε*] is considered an irreducible error. For simplicity, we define the prediction error as *r* − *v*(*s*), although there are variants, such as *r* + *γv*(*s* ′) − *v*(*s*), *r* + *γ*max_*a*′_*v*(*s* ′,*a* ′) − *v*(*s*). Accordingly, the reward maximization problem can be written as 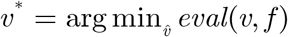, which we call *value learning* (Figure 1B).

We consider two standard strategies for value learning: MB and MF learning. The computational complexity of MB is greater than that of MF. It follows that the bias error of the MB learning agent is usually smaller than the bias error using MF, whereas the variance error of MB learning is greater than that of MF learning, as illustrated in Figure 1A. These contrasts make it difficult to resolve the bias– variance tradeoff (Filipowicz et al., 2020), providing a strong motivation for combining MB and MF learning.

Accumulating evidence supports the idea that the brain combines these two types of learning (Daw et al., 2005, 2011; Gläscher et al., 2010), e.g., *v*(*s*;*w*) = *wv*_MB_(*s*) + (1 − *w*)*v*_MF_(*s*). as previously discussed, the bias-variance characteristics of MB and MF learning are not linearly dependent; therefore, for a given environment (*f*, *ε*), there exists a minimizer *w*\* for (1) such that (*w*\*,*v*\*) = arg min_*w,v*_ *eval*(*v*, *f*), which can be achieved through value learning. Accordingly, the lower bound of (1) depends on the type of learning strategy used by the agent.

### Strategy control for bias-variance lower-bound minimization

In a situation in which the environmental context changes (*f* → *f* ′), the above minimum condition associated with the minimizer *w*\* may no longer hold, that is, the lower bound of the evaluation may increase:

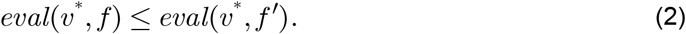

This provides two important implications. First, tracking the evaluation function value, that is, the average amount of prediction error, can serve as a reliable indicator of the context change. Second, to cope with changes in context, the agent needs to readjust the balance between the model-based and model-free learning strategy, i.e., decrease the lower bound of the evaluation in (1). This can be achieved by solving *w*′ = argmin_*w*_ *eval*(*v, f*′) while minimizing (1) through value learning. We call this a *strategy control* (Figure 1B; See Methods).

Previous studies provide neural evidence showing that the lateral prefrontal cortex implements a dynamic arbitration between model-based and model-free learning (adjusting *w*), which is based on a prediction error-based reliability computation (*eval*(*v*, *f*)) (Kim et al., 2019; Lee et al., 2014; Weissengruber et al., 2019).

### Adaptive control is a near-optimal solution to lower bound minimization

Although dynamic arbitration between MB and MF learning provides a means to decrease the lower bound of the bias– variance tradeoff to a great extent (strategy control), this does not consider the effect of the irreducible noise accompanied by context change on the evaluation function. To accommodate this situation, the change in the environmental context should be specified as (*f*, *ε*) → (*f* ′, *ε*′), as opposed to *f* → *f* ′. Under this new assumption, condition (2) reflects not only changes in the bias-variance, but also changes in irreducible noise variance var[*ε*], making it difficult to estimate the lower bound of the bias-variance. In other words, the noise variance arising from the environmental context complicates the minimization process of the bias-variance lower bound. To address this issue, the agent needs to adapt to the changing noise conditions, which we call *adaptive control* (Figure 1B).

One simple remedy for this issue is a baseline correction, for which the agent tracks the changes in the variable that imposes variability on its estimation to factor it out of the estimation. Although this idea has been successfully tested in machine learning (Degris et al., 2012; Kim et al., 2018; Ng et al., 1999; Wang et al., 2016), no neural evidence has been made available. Finally, by applying the baseline correction for the prediction error to the strategy control problem, our evaluation problem boils down to a simple bias-variance estimation.

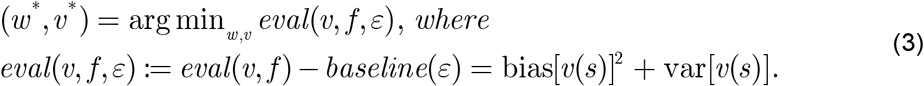

In summary, an ideal agent minimizes the amount of prediction error (*value learning*) by combining the model-based and model-free learning strategies based on the predicted performance (*strategy control*), and this ability is further facilitated by the baseline correction for prediction error (*adaptive control*). Note that the implementation relies on a single variable, i.e., the prediction error.

### Markov decision task with context changes

A test of our theory requires a learning task with changes in context. For this, we used a two-stage Markov decision task (Lee et al., 2014) with two context variables: a reward goal (observable task context) and state-transition uncertainty (latent task context) (see Figure 2A). The dataset consists of behavioral data on 60 subjects that we collected for this study and fMRI data on 22 subjects from a previous study (Lee et al., 2014), which amounted to a total of 82 behavioral data and 22 fMRI data (38 women, aged 19–40 years).

**Figure 2.**
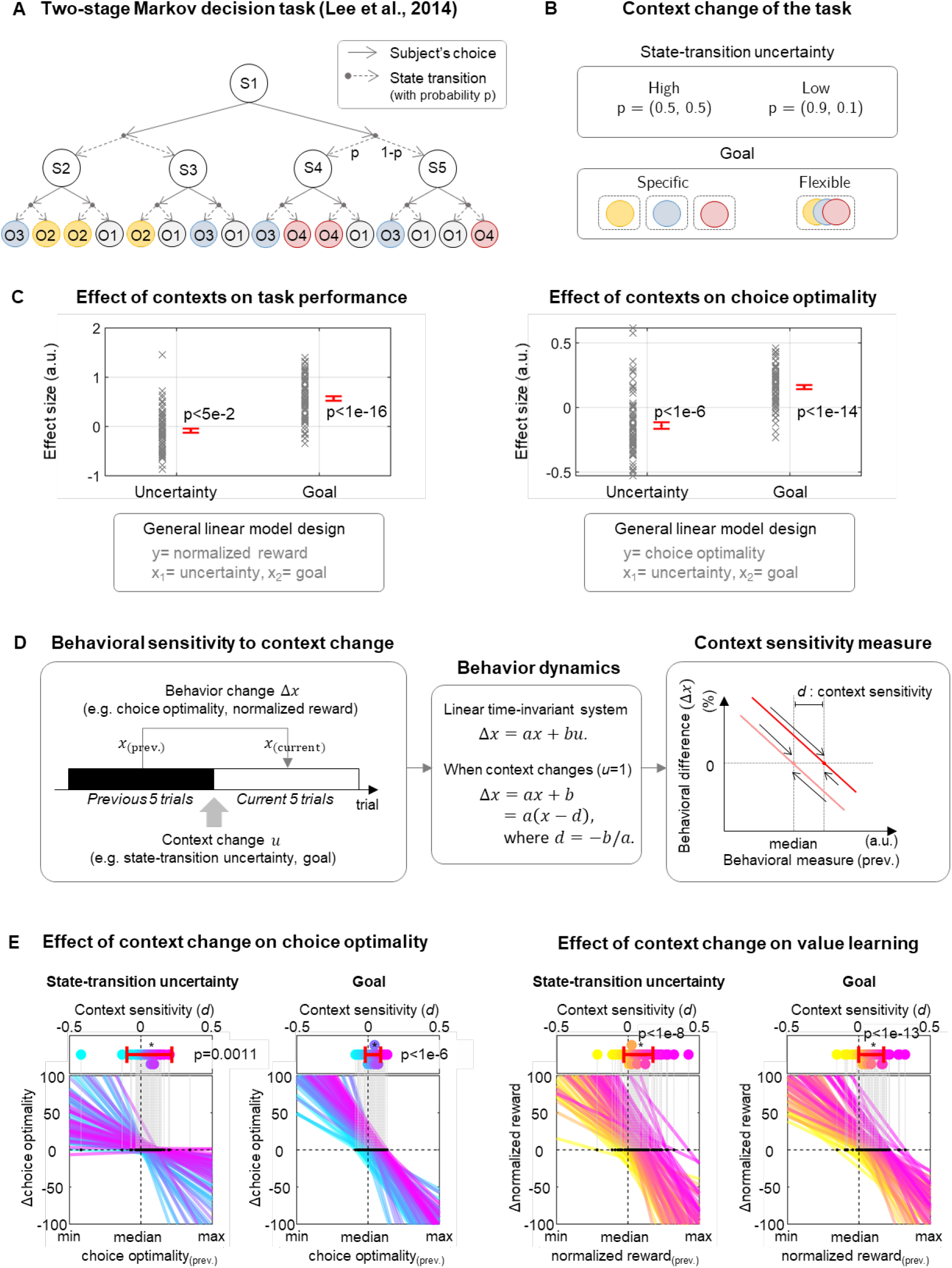
Evidence of context-sensitive behavioral control. **(A)** We collected data using a two-stage Markov decision task (Lee et al., 2014) (N = 82; among which, data on 22 subjects were from the original study). Each circle and arrow represent a state and action, respectively. **(B)** Two types of context were manipulated during the task: state-transition uncertainty and the reward goal. **(C)** Effect of contexts on behavior (general linear model analysis). Both an uncertainty and reward goal have significant effects on the choice optimality (paired t-test; t_81_ = 5.5570 and t_81_ = 9.8581, p < 1e68 and p < 1e-14, respectively), and normalized reward (paired t-test; t_81_ = 2.0153 and t_81_ = 12.4788, p < 5e-2 and p < 1e-16, respectively). Error bars indicate the SEM across subjects. For full details, see Supplementary Figure 3 and Table 1. **(D)** Behavioral measure to quantify the sensitivity to a change in context. We measured the degree of behavior changes (Δx) caused by context changes (*u*) (left panel). The measure is specified as a linear dynamical system, accommodating both the Markovian property of, and the effect of context change on, the behavior (the middle panel). Finally, the context sensitivity measure (*d*) is defined as the degree of context change effect (*b* in the middle panel) relative to the inherent behavioral dependency (*a* in the middle panel). For more details, see Methods. **(E)** The context change (state-transition uncertainty and goal) influences subjects’ performance, i.e., choice optimality (left) and normalized reward (right). In all four cases, the effect sizes were highly significant (paired t-test; for the context effect on the choice optimality, t_79_ = 6.7472 and t_81_ = 4.9393, with p = 2.2650 × 10^−9^ and p = 4.1438 × 10^−6^, respectively, and for the normalized reward, t_81_ = 6.3885 and t_81_ = 9.0438, with p = 9.9323 × 10^−9^ and p = 6.5170 ×10^−14^, respectively; we disregarded certain outliers with fitted lines with overly small and positive slope values because it means there is no behavioral control owing to a change in context). Error bars indicate the standard deviation (SD) across subjects.

This task involves two types of contexts (Figure 2B). The first context, the reward goal, was manipulated within the task. It included two types of conditions regarding the reward goal: specific and flexible. Under the condition of the specific goal, at the start of each trial, the participants were informed of the color of the coin redeemable for that trial. Each colored coin was associated with a different reward value (40, 20, and 10). The redeemable coin color was switched on a trial-by-trial basis. Under the condition of the flexible goal, all colored coins were redeemable. Because the goal of this task was to maximize the total amount of reward, it was reasonable to expect that the participants were motivated to collect coins with a maximum value of 40 during this condition. The context of the second task, the state-transition uncertainty, is switched between low (state-action-state transition probability p = [0.9, 0.1]) and high (p = [0.5, 0.5]). Changes in state-transition uncertainty were implemented to induce a low or high SPE, thereby encouraging the MB or MF RL, respectively. Note that the state-transition probability changes were implicit, and thus the participants were expected to learn from experiencing multiple episodes of state-action-state transitions (see Supplementary Figure 1 for full details of the relationship between task context and optimal RL). All participants conducted the task successfully (hit rate > chance level; paired t-test; t_81_ = 13.4226, p = 1.3555 × 10^−22^).

### Behavioral evidence of adaptive control

To assess the subjects’ ability of value learning and strategy control, we used two behavioral measures: normalized reward and choice optimality. The former, the equivalent of a hit rate, is a well-known behavioral indicator of value learning, whereas the latter is known as a key behavioral indicator of model-based control (Kim et al., 2019). The normalized reward is defined as the obtained reward value divided by the maximum amount of reward in each trial, which is constrained by redeemable coin color (see Methods). The choice optimality, is defined as the proportion of trials in which the subjects’ choice to be the same as that of the ideal agent, assuming the ideal agent making choices based on full information about the current contexts. (see Methods). Note that the choice optimality was previously shown to be more reliable than the other behavioral measures, such as choice consistency (Kim et al., 2019) (Supplementary Figure 2). Both contexts, goal and state-transition uncertainty, had a significant effect on normalized reward and choice optimality (Figure 2C; general linear model analysis, p < 1e-5), providing behavioral evidence of value learning and strategy control (the first and second components of Figure 1B).

In a subsequent analysis, we deliberately exploit these two behavioral measures to explore behavioral evidence of adaptive control (the third component of Figure 1B). One defining characteristic of adaptive control is that a change in control input (choice behavior) is influenced not only by its previous behavior, but also by changes in the environment (context change). To fully accommodate this situation, we modeled each choice as a linear dynamical system with a context input as an intervention (Figure 2D; see Methods). Note that this approach allows us to separate out the effect of a change in context (“*b*” in Figure 2D) from the behavioral variability arising from value learning (“a” in Figure 2D), which is more efficient than a simple regression or general linear model. As a result, the effect of context changes on the choice behavior, called “context sensitivity,” is defined as the ratio of context change effect to inherent behavioral dependency (see Methods).

We found a highly significant effect of context changes (both state-transition uncertainty and goal) on both choice optimality and normalized reward (Figure 2E, p<1e-3), implying that subjects incorporate changes of context into both strategy control and value learning. These results provide behavioral evidence of adaptive control.

### Adaptive control of model-based and model-free learning

The theoretical idea of the bias–variance tradeoff in the context of reinforcement learning motivates us to implement a computational model of adaptive control (Figure 3). The model consists of three functional components. First, the two types of value learning, model-based (MB) and model-free (MF) RL, update state-action values by applying a state prediction error (SPE) and a reward prediction error (RPE), respectively (indicated by the red and blue light lines in Figure 3A). This corresponds to the value learning part shown in Figure 1B.

**Figure 3.**
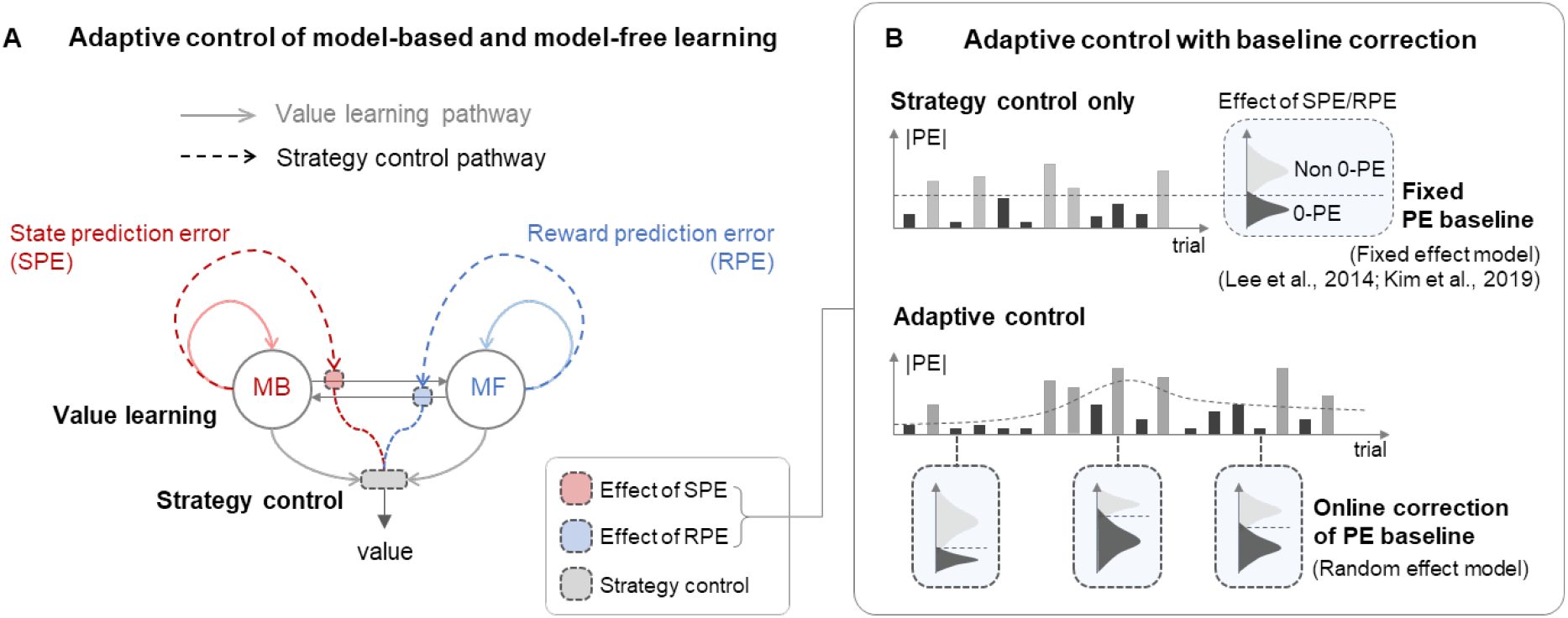
Computational model of adaptive control (implementation of Figure 1B with model-based and model-free RL). **(A)** Schematic diagram of adaptive control of model-based and model-free RL. Light color lines and dotted lines indicate the value learning and strategy control pathway, respectively. This concerns two types of value learning strategies, model-based (MB) and model-free (MF) RL, each of which uses state prediction error (SPE) and reward prediction error (RPE), respectively. **(B)** Adaptive control. A pure strategy control is based on binary classification of zero and non-zero PE distributions with a fixed baseline (top; fixed effect model). Adaptive strategy control has an ability to correct for prediction error changes (bottom; random effect model).

Second, strategy control is built on top of value learning. This corresponds to the strategy control part shown in Figure 1B. To measure the success in making predictions with the current value learning strategy, the SPE and RPE are converted into the prediction reliability of MB and MF RL, respectively (indicated by the red and blue dotted lines in Figure 3A). Here, the prediction reliability is a function of the number of times that the amount of prediction error exceeds a prediction error baseline (the top of Figure 3B) (Kim et al., 2019; Lee et al., 2014).

Next, we consider another version of strategy control, called adaptive control (Figure 3B). This corresponds to the adaptive control part shown in Figure 1B. Unlike the strategy control with a fixed prediction error baseline, the adaptive control continually updates the baseline to fully accommodate the changing level of irreducible prediction error induced by context changes (the bottom of Figure 3B).

The implementation is based on a Dirichlet process Gaussian mixture model (DPGMM), the nonparametric Bayes approach known to be effective in discovering skills or clustering different sizes of actions in a reinforcement learning task (Niekum and Barto, 2011).

Our adaptive control scheme was compared with a pure strategy control. In the pure strategy control, the computation of the prediction reliability is based on the binary classification of zero and non-zero prediction errors with a fixed threshold (Kim et al., 2019; Lee et al., 2014). This means that a reliability evaluation may be impaired when the agent is placed in other contexts with a different level of irreducible prediction error. By contrast, the adaptive control learns and updates the zero and non-zero prediction error distributions by updating the posterior distributions of the prediction error clusters on a trial-by-trial basis (see Supplementary Section A), making it possible for the agent to reliably measure the success of the current value learning strategy while automatically correcting for the long-term average of the prediction error. Note that this will help resolve the bias–variance tradeoff issue illustrated in Eq. (3).

### Model-based evaluation of multiple competing hypotheses

To examine whether our adaptive control hypothesis explains the choice behavior of the subjects, we ran two-fold model comparison analyses: a Bayesian model comparison (Figure 4A) and a parameter recovery test (Figure 4B). The former is intended to assess underfitting; it quantifying the extent to which a model accounts for the behavioral data of the subjects after correcting for model complexity. The latter is intended to test whether the model suffers from an overfitting.

**Figure 4.**
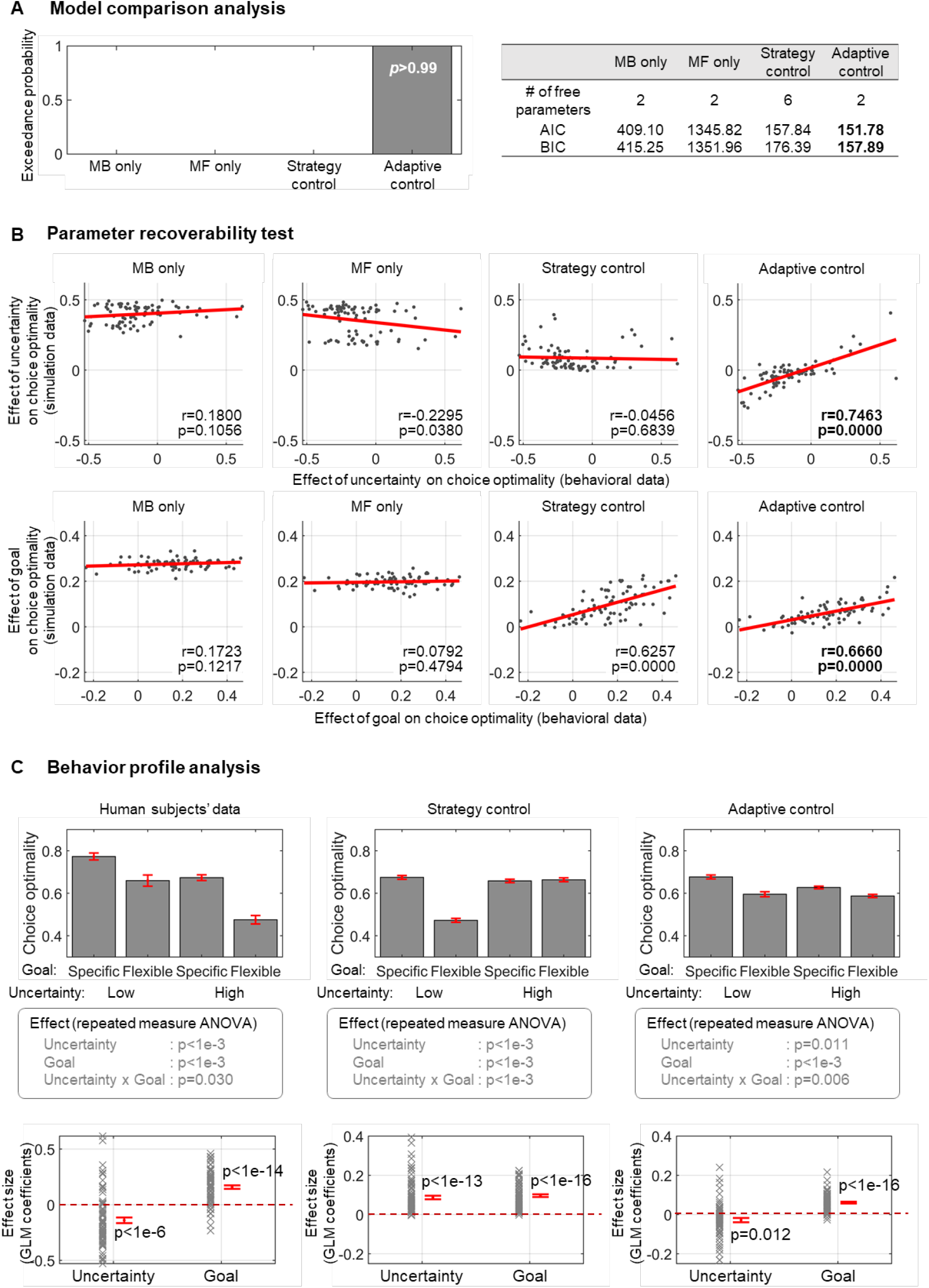
Model comparison analyses. **(A)** Bayesian model comparison analysis. We fitted each model to the data of each subject to compute the likelihood of choices, and then ran Bayesian model comparison analyses to correct for model complexity. Notably, the exceedance probability is larger than 0.99, strongly supporting the adaptive control hypothesis. Shown on the right are the results with other performance metrics (AIC and BIC). **(B)** Parameter recovery analysis (overfitting test). For this, we measured the effect of the task contexts on the choice behavior for the behavioral data of the original subjects and the simulation data generated by the model fitted to the original data and compared them. The task context effect was computed using the GLM analysis. To ensure the reliability of the simulation, we used the simulation data from 40,000 trials for each subject. **(C)** Behavioral profile analysis to assess the effect of task contexts on the choice behavior. The first row shows comparison of choice optimality between human subjects’ data and the top 2 models (strategy control and adaptive control) from the above two analyses (A) and (B). The second row is the results of 2-way repeated measures ANOVA (also see Supplementary Table 3 for full details of the adaptive control model). The third row shows the effect of contexts on choice optimality, computed using a general linear model analysis (GLM). The GLM results on the human subjects’ data were taken from Figure 2C. Error bars indicate the SEM across subjects.

We compared the performance of six different models in terms of the likelihood of choices (see Supplementary Figure 4 and Supplementary Table 2 for a full description of the six models). They include two value learning models (MB and MF), a strategy control model (Lee et al., 2014), and three different versions of adaptive control. The first two versions are equipped with adaptive control for either SPE or RPE. The third version is a full adaptive control model for both SPE and RPE (Figure 3A). Figure 4A summarizes the performance comparison between the four main models.

In the model comparison analyses, we used three different performance metrics: Bayesian model selection (Stephan et al., 2009) (BMS), Bayesian information criterion (BIC), and Akaike information criterion (AIC). We found that the full adaptive control model outperformed all other models in all performance measures with an extremely high confidence level (Figure 4A; see Supplementary Figure 4A for full comparison). When we compared only the top three models, this model was significantly better than the others (Supplementary Figure 4B; paired t-test, *p* < 1e-15).

To preclude the possibility of an overfitting, we ran parameter recovery tests (Figure 4B; see Methods, parameter recovery analysis). For each model, we compared the task context effects on the choice behavior in the behavioral data of the original subjects with those in the simulation data generated by the model fitted to the original data. Notably, the adaptive control model is the only version showing a significant positive correlation between the behavioral and simulation data. This also means that some behavioral features associated with value learning falsify alternative models, including value learning and strategy control.

To more directly demonstrate the accountability of the model for the trial-by-trial choice behavior of the subjects (left action L versus right action R), we compared the proportion of the right action of the subjects (action R) with the model-predicted probability of choosing the right action (Supplementary Figure 5). When measuring the choice optimality as a function of task contexts, we found that the adaptive control is the only version whose behavior profile matches that of human subjects (the top row of Figure 4C). This is confirmed by the GLM analysis assessing the context effect on the choice optimality (the bottom row of Figure 4C). In summary, all model evaluation results strongly support our adaptive control hypothesis.

### Role of prediction error baseline in adaptive control

To understand the nature of adaptive control, we investigated the role of prediction error in context-sensitive adaptive control. Our analyses are intended to show that the baseline estimation of the prediction error reflects changes in context (Figure 5A-C), which essentially leads to a behavioral adaptation (Figure 5D).

**Figure 5.**
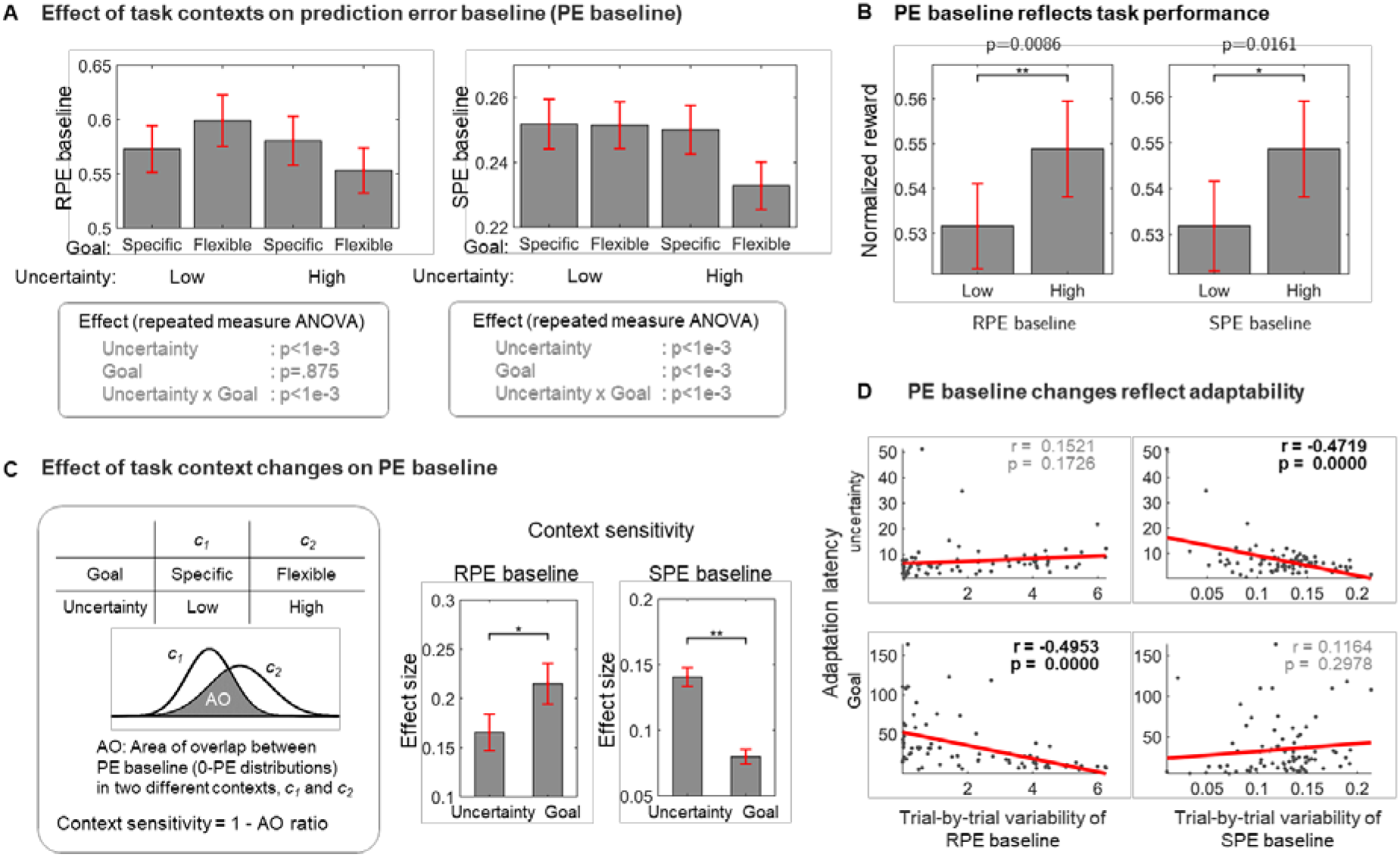
Baseline estimation of prediction error of the adaptive control model reflecting task context information. (A) Effect of task contexts on prediction error (PE) baseline. The PE baseline is defined as the mean of zero PE to track its updating trends. The results of two-way repeated measures ANOVA are shown at the bottom. (B) Relationship between the PE baseline and task performance (normalized reward). (C) Effect of task context changes on PE baseline. The context sensitivity was defined as the area of overlap between two PE baseline distributions in each task context (left). The RPE and SPE baseline is highly sensitive to goal changes (paired t-test; t_81_ = 2.6311, p = 0.0102) and uncertainty changes (paired t-test; t_81_ = 5.9969, p = 5.336 × 10^−8^), respectively. (D) Relationship between the amount of PE baseline changes (trial-by-trial variability of the PE baseline) and adaptation performance (adaptation latency). The adaptation latency (y-axis) is defined as the number of trials that it takes for the model to begin using an optimal learning strategy following the context change. The optimal strategy is defined as the strategy the ideal agent uses in each context. The ideal agent is assumed to make choices based on full information about the current contexts, leading to the optimal task performance. The optimal strategies for the specific goal, flexible goal, low uncertainty, and high uncertainty condition are model-based, model-free, model-based, model-free learning, respectively. The trial-by-trial variability of the PE baseline (x-axis) quantifies the amount of the PE baseline updates. Error bars are SEM across subjects.

We determined the interaction effect of the contexts (state-transition uncertainty and goal change) on the baseline estimation of the prediction error (Figure 5A). We also found that the PE baselines reflect the task performance (Figure 5B). These results, while providing circumstantial evidence that the PE baseline encodes task contexts, demand a close investigation into how the PE baseline responds to changes in context.

Thus, we measured the sensitivity of the PE baseline to changes in context. The context sensitivity was defined as the amount of change in the baseline prediction error distribution induced by changes in context (the left gray box of Figure 5C). Interestingly, we found a dissociable effect of the contexts on the RPE and SPE (see the right side of Figure 5C; two-way repeated measures ANOVA, p < 1e-3). Specifically, the RPE baseline distribution is more sensitive to changes in goal than that of SPE, and the opposite effect was found for the case of the SPE baseline. This is consistent with the fact that the RPE and SPE reflect discrepancies in predictions regarding the value and state transition, respectively.

To further examine the role of the PE baseline in adaptation, we assessed the adaptability of the model as a function of the amount of PE baseline changes (Figure 5D). To do so, we calculated the latency of switching to an optimal strategy that an ideal agent uses following changes in context. The lower the latency, the better the adaptability. Here, the ideal agent is a hypothetical agent making optimal choices based on full information regarding the current contexts (goal and state-transition probability values) (Kim et al., 2019). The results showed that the amount of change in the SPE baseline becomes negatively correlated with the adaptation latency after changes in the state-transition uncertainty (*R*^*2*^ = 0.223; correlation coefficient *r* = −0.472, *p* < 1e-4), whereas the RPE baseline change is negatively correlated with adaptation latency following goal changes (*R*^*2*^ = 0.245; correlation coefficient *r* = −0.495, *p* < 1e-4). Taken together, our results corroborate the view that context-sensitive learning leads to a fast adaptation.

### Neural computations underlying adaptive control in the prefrontal cortex

We intended to provide a neural level account for adaptive control by regressing the key variables of our model against the fMRI data (see Supplementary Table 4 for voxel-level statistics). Our neural analyses consisted of three steps: identifying neural representations of the key variables for model-free and model-based learning (corresponding to *value learning* of Figure 1B), finding a key variable for allocating control over these two types of learning (corresponding to *strategy control* of Figure 1B), testing the neural effect of context-sensitive strategy control (corresponding to *adaptive control* of Figure 1B).

The first step of our fMRI analysis explores neural evidence of the two key variables sufficient for *value learning*: prediction error and action value. Specifically, we found evidence of state prediction error of model-based learning in the dorsolateral prefrontal cortex (dlPFC) and bilateral insula (one-sided t-test; t_21_ = 8.36 for left dlPFC, t_21_ = 11.09, and t_21_ = 11.12 for left/right insula; and peak-level family wise error (FWE) corrected, p < 0.05) as well as evidence of reward prediction error of model-free learning in the dorsal and ventral striatum (t_21_ = 6.96 for the right ventral striatum; peak-level FWE corrected at p < 0.05; t_21_ = 5.33 for right ventral striatum and t_21_ = 5.41 for dorsal striatum; and cluster-level FWE corrected p < 0.05), all of which are fully consistent with previous findings (Gläscher et al., 2010; Lee et al., 2014; McClure et al., 2003; O’Doherty et al., 2003). Note that owing to our task design for separating our model-based from model-free learning, these two types of prediction error signals are not highly correlated (average *r*=−0.2854). Our neural results for value signals are also highly consistent with those of previous studies. The value signal for model-free learning was correlated with neural activity in multiple brain regions, including the supplementary motor area (SMA), dlPFC, anterior-lateral PFC (alPFC) (Hare et al., 2011; Rowe et al., 2010) (t_21_ = 6.56 for SMA; peak-level FWE corrected, p < 0.05; t_21_ = 5.28 for dlPFC and t_21_ = 3.96 for alPFC; cluster-level FWE corrected at p < 0.05), and posterior putamen (Tricomi et al., 2009; Wunderlich et al., 2012) (t_21_ = 3.99; after a small volume correction [SVC], peak-level FWE corrected p < 0.05). Evidence of the value signal for model-based learning was found in the orbital and medial PFC (omPFC) (Lee et al., 2014) (t_21_ = 3.60; after SVC, peak-level FWE corrected, p < 0.05). Finally, the integrated value, given by a combination of model-free and model-based value signals, was found in the ventromedial PFC (vmPFC) (Boorman et al., 2009; Hare et al., 2009; Lee et al., 2014; Rushworth et al., 2011) (t_21_ = 4.77 for vmPFC; cluster-level FWE corrected, p < 0.05).

The second step investigates neural correlates for *strategy control*, for which we examined the evidence of arbitration between model-based and model-free value learning. The key variable for arbitration is the reliability signal (Kim et al., 2019; Lee et al., 2014), a prediction error-based proxy for the predicted performance of each value learning strategy. Critically, we found that reliability information was reflected in the neural activity patterns in the ventrolateral PFC (vlPFC) bilaterally and frontopolar cortex (FPC) (Kim et al., 2019; Lee et al., 2014) (t_21_ = 6.17 for left vlPFC, t_21_ = 5.60 for right vlPFC, t_21_ = 5.22 for FPC; cluster-level FWE corrected, p < 0.05). Taken together, using our new model, we successfully replicated all previous findings supporting the existence of value learning and strategy control.

The last step is focused on investigating a neural computation underlying *adaptive control*, for which we explored the neural correlates of the prediction error baseline, the key variable for such control (related to Figures 3B and 6), followed by testing its effect on the brain areas encoding the key variables for both *strategy control* and *value learning*. We found that the SPE baseline is reflected in the neural activity of the fusiform gyrus (FFG) and lingual gyrus (LG) (t_21_ = 5.32 for FFG, t_21_ = 4.04 for LG; cluster-level FWE corrected, p < 0.05). The RPE baseline effect is shown in the pre-supplementary motor area (pre-SMA) (Supplementary Figure 6A; t_21_ = 7.23; peak-level FWE corrected, p <0.05; see Supplementary Section B for a full analysis and Table 4 for voxel-wise statistics). Note that these two types of baseline prediction error signals are not highly correlated (average *r*=0.0592).

**Figure 6.**
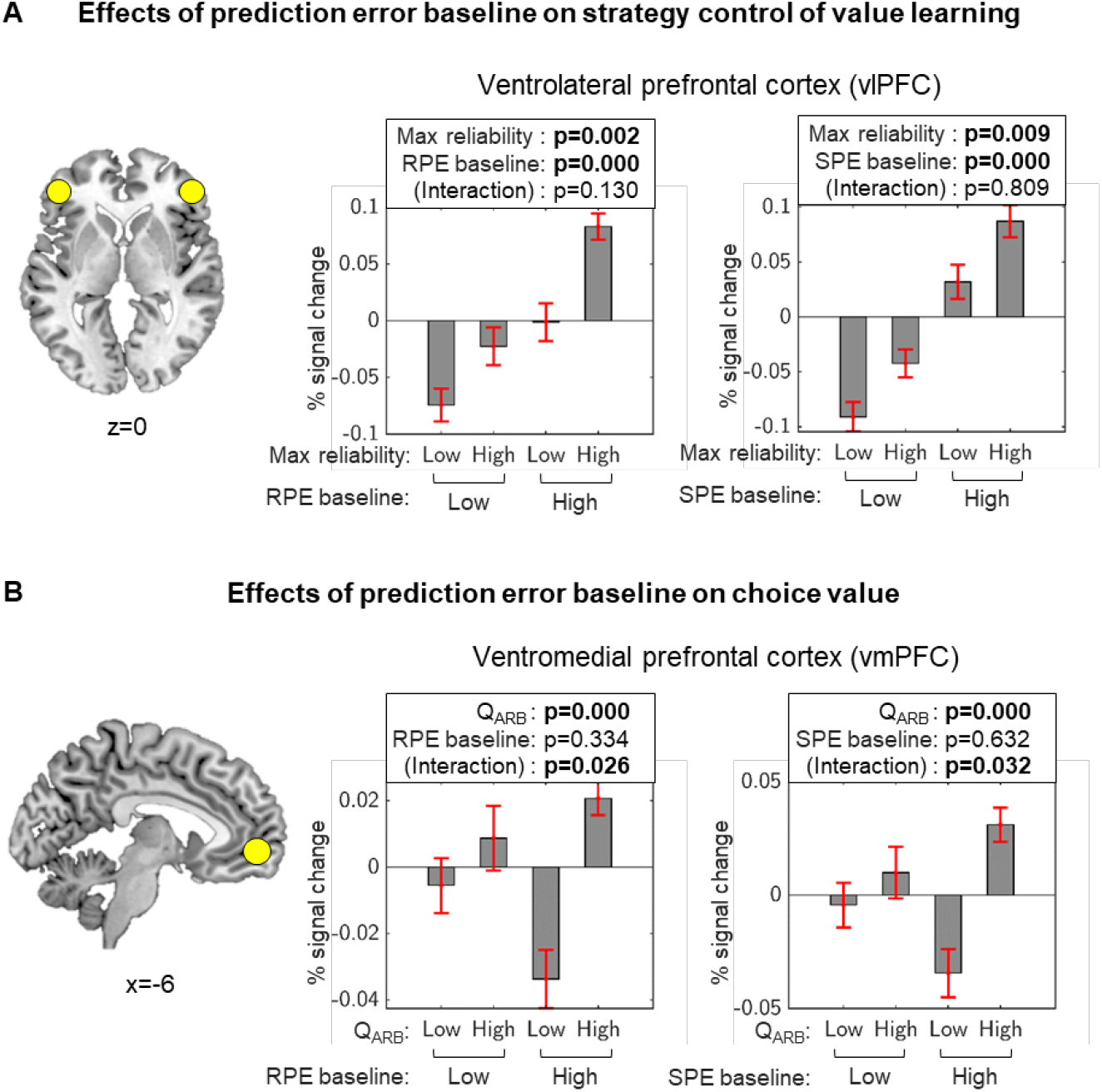
Neural evidence of adaptive control. **(A)** Percent of signal changes in the bilateral vlPFC as a function of the Max reliability of value learning strategies (model-based and model-free) and of the prediction error baseline. **(B)** Percent of signal changes in vmPFC as a function of the chosen value and of the prediction error baseline. The chosen value is given by *Q* (*s*,*a*) =*wv*_MB_ (*s*,*a*) +(1 − *w*)*v*_MF_ (*s*,*a*), where *s, a* is each state and the action taken at the state *s*, respectively; *v* refers to the value computed from the model-based (MB) and model-free (MF) strategy; *w* is the model choice probability computed by the strategy control. Data are split into two equal-sized bins according to the 50th percentile. Results of two-way repeated measures ANOVA are shown with the error bar plots. Error bars are SEM across subjects.

Critically, we found the PE baseline effect on both the vlPFC and vmPFC, the two key regions for strategy control and the corresponding value integration. The neural activity of the vlPFC significantly increased with the prediction error baseline (Figure 6A; main effect, *p* < 1e-3), which is fully consistent with the view that the predicted performance reliability of the current value learning strategy (max reliability) increases when the agent makes largely correct predictions (increasing the PE baseline). The increase in the PE baseline significantly improved the value representation in the vmPFC (Figure 6B; interaction effect, *p* < 0.05). These findings show that the vlPFC and vmPFC implement computations necessary for adaptive control.

### Effect of adaptive control on prefrontal-striatal functional connectivity

Following our findings showing that the key variable for adaptive control (prediction error baseline signal) is reflected in the neural activity in vlPFC and vmPFC, we ran functional coupling analysis to better understand the effect of adaptive control on the strategy control - model-free learning – valuation network (Figure 7A). For this we consider the brain area for strategy control (bilateral vlPFC; Lee 2014; Kim 2019), model-free learning (left posterior putamen; Lee 2014; Tricomi 2009; Wunderlich et al., 2012; Kim 2019), and valuation (vmPFC; Boorman et al., 2009; Hare et al., 2009; Lee et al., 2014; Rushworth et al., 2011) (Figure 7B).

**Figure 7.**
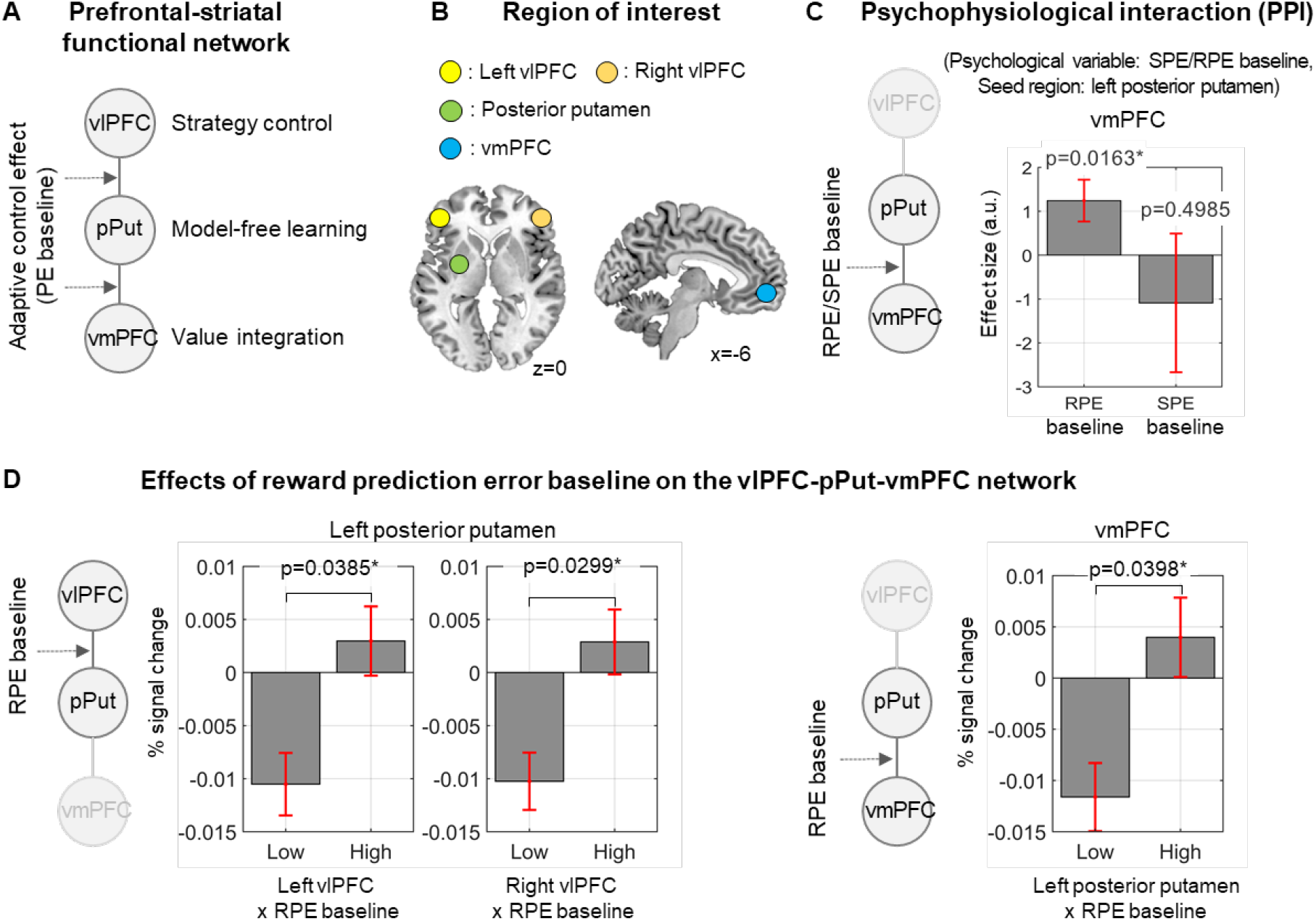
Adaptive control effect on the vlPFC-posterior putamen-vmPFC network. **(A)** Prefrontal-striatal functional network for value-guided learning. This network involves the brain areas for strategy control (bilateral vlPFC; Lee 2014; Kim 2019), model-free learning (left posterior putamen; Lee 2014; Tricomi 2009; Wunderlich et al., 2012), and valuation (vmPFC; Boorman et al., 2009; Hare et al., 2009; Lee et al., 2014; Rushworth et al., 2011). **(B)** The corresponding regions of interest for the network analysis. **(C)** Psychophysiological Interaction (PPI) analysis to assess the parametric modulation effect of prediction error baseline on model-free valuation. Based on the prior reports (Lee 2014; Tricomi 2009; Wunderlich et al., 2012), we focus on the interaction between left posterior putamen and vmPFC. **(D)** Effect of reward prediction error baseline on the vlPFC-posterior putamen-vmPFC network. The neural activity level (percent signal change) is shown as a function of the two different trial types: one in which an interaction value (seed region’s neural activity x RPE baseline) is low and the other associated with high value.

We found in the psychophysiological interaction (PPI) analysis that valuation of model-free learning is parametrically modulated by the RPE baseline (mean effect size = 1.24, paired t-test; t21 = 2.61, p = 0.0163), but not SPE baseline (mean effect size = −1.09, paired t-test; t21 = 0.69, p = 0.4958) (Figure 7C; For full network analysis, see Supplementary Figure 7 and Supplementary Section C). This result is fully consistent with previous view that the striatal model-free system guides valuation (Wunderlich et al., 2012; Lee 2014). It is further corroborated by the direct effect of RPE baseline on the downstream of the vlPFC-pPut-vmPFC network: between vlPFC and posterior putamen (the left of Figure 7D; paired t-test t_21_ = 2.2075 p = 0.0385 for the left vlPFC, t_21_ = 2.3292 p = 0.0299 for the right vlPFC) and between posterior putamen and vmPFC (the right of Figure 7D; paired t-test t_21_ = 2.1913 p = 0.0398).

## Discussion

In the present study, the hypothesis that the brain has the capacity to resolve the bias–variance error minimization problem was formally tested. For this, we presented a theoretical idea that adaptive control of model-based and model-free reinforcement learning can resolve a tradeoff between bias and variance error. Based on this framework, we proposed a simple and near-optimal solution in the presence of changes in the task context. The key component for resolving this problem is the baseline correction for prediction error, which serves to reduce the lower bound of the bias–variance error by factoring out an irreducible noise in the prediction error. We used data from 82 subjects collected with a two-stage Markov decision task in which the two types of changes in context, i.e., goal and state-transition uncertainty, and showed behavioral evidence of adaptive control. To formally test this idea, we implemented a computational model that dynamically combines model-based and model-free learning (strategy control) strategies while correcting for the prediction error baseline (adaptive control). Using this model, we were able to elucidate the role of the prediction error baseline in encoding task context changes and further provide a detailed account of how doing so helps improve the context adaptability. In the subsequent model-based fMRI analysis with data from 22 subjects, we replicated the previous findings of value learning and strategy control. As the key finding of the neural analysis, neural activity patterns of the ventrolaternal and ventromedial prefrontal cortex, i.e., the brain locus previously known to guide the strategy control and value learning, respectively, reflect the changes in the prediction error baseline. This result suggests multiplexed representations of the prefrontal cortex to improve the adaptability of the strategy control of model-based and model-free learning.

Our study has a few significant implications for artificial intelligence. First, our approach lays the necessary groundwork for understanding the brain’s ability to reliably learn complex tasks despite changes in context. This can directly stimulate various task-independent and generalizable algorithm design, such as context-aware reinforcement learning and meta-learning. Second, our behavioral, computational, and neural account for how the prefrontal cortex minimizes the bias and variance prediction error offers a valuable insight into the development of learning systems that maintains a balance between performance and efficiency (Lee et al., 2019). For example, human’s preference for model-free learning helps reducing reaction time and cognitive loads while potentially compromising performance. On the other hand, model-based learning can improve task performance and sample efficiency though it requires a relatively large amount of cognitive resource. According to our theoretical view, whilst the strategy control can deal with this tradeoff, its efficiency is reduced by context changes. Our results (Figure 3, 5 and 6) provide a conceptual solution to this issue: multiplexed latent representations of predicted performance reliability and baseline prediction error.

Our proposition advanced the conventional view of the prediction error, arguing that the prediction error represents not only the discrepancy between an actual outcome and what the agent expects to obtain, but also the least and most of what the agent expects to obtain from the environment. The primary role of the prediction error is to quantify the amount of information that the agent wants to acquire, providing the need to reinforce the current value learning policy. The secondary role is to estimate the irreducible amount of prediction error contingent on the task contexts, facilitating the context-dependent allocation of behavioral control to different value learning strategies. One interesting interpretation would be that when the agent finds itself in a novel situation owing to a change in the latent task structure, the agent seems to lower the expectation of its own performance, and therefore would not be surprised by a certain amount of prediction error.

This view is corroborated by a model-based analysis. The computational model implementing our hypothesis was shown to account for the behavior patterns of the subjects significantly better than alternative hypotheses, including pure value learning (model-based or model-free only) and strategy control (Lee et al., 2014). Note that the prediction error baseline estimation of our model accommodates a highly plausible and realistic scenario. During each trial, the model classifies the prediction error into two cases: zero and non-zero prediction errors, simply representing correct and incorrect, respectively. Classification and updates of the two prediction error distributions are performed purely online (see Methods). We used the SPE and RPE, each of which we hypothesized as being sensitive to different types of changes in context, such as reward goal and state-transition uncertainty (Gläscher et al., 2010; Kim et al., 2019; Lee et al., 2014).

Although using the prediction error baseline is perhaps the simplest way to estimate the irreducible error constrained by the given context, there may exist other variables that better accommodate the changes in context. For example, the proportion of unrewarded trials affects the activation patterns of dopamine neurons in subsequent trials (Nakahara et al., 2004), and both unexpected and expected rewards affect the response of dopamine neurons in the ventral tegmental area (Eshel et al., 2016). The neural correlates of unexpected uncertainty found in multiple cortical regions (Payzan-LeNestour et al., 2013) suggest the possibility of a context sensitivity of an SPE. Recent findings suggest that unsigned and signed RPEs separately contribute to the agent’s value update (Haarsma et al., 2018; Iigaya et al., 2020). Future studies should attempt to elucidate the role of these variables in adaptive control.

Our neural findings (Supplementary Figure 6) indicate that the prediction error baseline reflects changes in context. As noted earlier, we found the SPE baseline effect in both the fusiform gyrus and lingual gyrus. This is broadly in agreement with the previous finding that the fusiform gyrus is engaged in context-dependent novelty processing (Biederman and Vessel, 2006; Habib et al., 2003; Henson et al., 1999; Stoppel et al., 2009). Moreover, our results implicating the dissociable sensitivity of the RPE and SPE baselines to explicit and implicit changes in context (Figure 5C and 5D) are consistent with the accumulating evidence that the neural activities in the fusiform gyrus reflect not only the novelty of the explicit context (e.g., visible cues) (Henson et al., 1999; Stoppel et al., 2009) but also the novelty of the implicit context (e.g., latent task structure). We also found a negative effect of the RPE baseline in the pre-SMA, the cortical area previously known for performance monitoring (Nee et al., 2011; Shenhav et al., 2013; Ullsperger et al., 2014). This corroborates our findings that the RPE baseline reflects the average task performance (Figure 5B and 5C).

To exclude alternative possibilities that may explain the behavioral and neural data, we examined the adaptive control hypothesis by systematically combining the following tests with stringent criteria: a Bayesian model comparison (Figure 4A), a context effect analysis based on the choice optimality measure (Figure 4B), a parameter recovery analysis (Figure 4C), and a Bayesian model selection on neural data in which different versions of the computational model were pitted against each other (Supplementary Figure 6B). Notably, all of the results consistently support our adaptive control hypothesis. The corresponding computational model provides a detailed account of how the brain minimizes a prediction error and effectively resolves the bias–variance tradeoff despite changes in context.

Finally, the neural results obtained using the computational model validate our theoretical claim about adaptive control (Figure 1B). As noted earlier, we found the effect of the key variable for adaptive control (prediction error baseline) on the brain area known to encode a value signal (vmPFC) as well as the brain area found to guide the strategy control of value learning (vlPFC). This result is further supported by a functional network analysis, in which we found that the functional connectivity between the vlPFC, vmPFC, and ventral striatum is regulated by the prediction error baseline (Figure 7). Taken together, our neural results elucidate how the brain incorporates changes in task context into value learning and its strategy control.

The implications of our findings can be assessed in broader contexts. Because the vlPFC, the functional loci for context-sensitive strategy control in our study, was previously implicated in related high-level cognitive functions, such as subjective confidence in decision making (De Martino et al., 2013), metacognitive processing, and uncertainty-driven learning rate control (O’Doherty et al., 2015), further investigation is warranted to explore the general role of the vlPFC as a meta-level cognitive controller. This view may be used to study complex traits of psychiatric disorders. Patients with obsessive-compulsive disorder (OCD), addiction, and related psychiatric disorders often show complex symptoms that cannot be explained through a single hypothesis. For instance, people with OCD exhibit highly habitual behaviors, supporting the idea of a bias toward model-free learning (Chamberlain et al., 2007; Gillan et al., 2011, 2014; Sjoerds et al., 2013; Voon et al., 2015), whereas others exhibit goal-directed behaviors that support model-based learning bias (Deckersbach et al., 2002; Joel et al., 2005; Rauch et al., 1997, 2007). These conflicting views can be reconciled by our adaptive control hypothesis, explaining the individual variability associated with a context-sensitive behavioral bias. In general, the corresponding theoretical idea may provide deeper insight into why a variety of high-level context changes, such as reward probability, state-transition probability, or volatility, are often implicated in psychiatric disorders.

## Acknowledgements

We would like to acknowledge our gratitude to John P. O’Doherty for approval of the use of the fMRI data. This study was supported by the National Research Foundation of Korea (NRF) grant funded by the Korean government (MSIT) (NRF-2019M3E5D2A01066267, Development of metacognitive AI for rapid learning; NRF-2017R1C1B2008972, Non-invasive control of human one-shot learning using brain-inspired AI), and Samsung Research Funding Center of Samsung Electronics under Project Number SRFC-TC1603-52.

## Author Contributions

Conceptualization, D. Kim and S. W. Lee; Methodology, D. Kim, and S. W. Lee; Software. D. Kim; Validation, D. Kim; Formal Analysis, D. Kim; Investigation, D. Kim, S. W. Lee; Resources, S. W. Lee; Data Curation, S. W. Lee; Writing – Original Draft, D. Kim; Writing – Review & Editing, J. Jeong and S. W. Lee; Visualization, D. Kim; Supervision, J. Jeong and S. W. Lee.

## Methods

A summary of the experimental methods relevant to the analyses described in the current study is provided here.

### Participants

A total of 82 adults participated in the current experiment, of whom 22 participants (i.e., 6 women, aged 19–40 years, with a mean age of 28 years) applied the task while being scanned using fMRI in the USA. This dataset was initially described in our previous report (Lee et al., 2014). The remaining 60 participants (31 women, aged 19–35 years, with a mean age of 22.75 years) underwent the experiment in South Korea. All participants were prescreened to exclude those with a history of neurological or psychiatric disorders. The studies were approved by the Institutional Review Board of the California Institute of Technology and Institutional Review Board of the Korea Advanced Institute of Science and Technology, respectively.

### Model-free (MF) and model-based (MB) reinforcement learning

To implement an MF learning agent, we used a SARSA, which uses conventional temporal-difference (TD) updates. The agent computes the reward prediction error (RPE) to update the state-action value. We also implemented the MB learning agent with FORWARD learning (Gläscher et al., 2010) and BACKWARD planning (Lee et al., 2014).The former allows an agent to learn about the model of the environment, whereas the latter allows instantaneous updates of the state-action value signal to reflect the change in goal given by the task. For each trial, during which a new goal was given, the model-based agent ran BACKWARD planning to perform instantaneous updates of the state-action value signal. Under each state, the agent used FORWARD learning, which computed the state prediction error (SPE) to update the state-action-state transition probabilities and corresponding state-action values.

### Behavioral measure

Choice optimality has been suggested as a behavioral measurement of MB control (Kim et al., 2019). This measures the proportion of trials in which the agent’s choice behavior to be the same as that of the ideal MB agent, who assumes to know the full information about the rewards and state-transition probability. Specifically, we added 1 if the agent and the ideal MB agent made a same choice, added 0 the agent’s choice was different from the ideal MB agent’s one, and added 0.5 when both choices are optimal due to the same expected rewards. However, unlike in the previous study (Kim et al., 2019), we strictly defined the amounts of reward in the outcome states. Thus, we normalized the rewards by dividing them with the maximum reachable reward (i.e., ranging from zero to 1). Specifically, for trials during the specific goal condition, except one, the other rewards are considered to be zero for computing the choice optimality.

### Degree of dependency on behaviors

We measured the degree of dependency on the behaviors based on the x-intercept from the first-degree polynomial fitting between behavioral measures in the previous block and its difference in the current block (Figure 2D). If the current behaviors do not depend on the previous behaviors, the behavioral measure in the previous block should be a median value when the behavioral difference is zero (pink line in Figure 2D). This indicates that subjects shift their behaviors regardless of how they performed in the previous block. By contrast, if the current behavior depends on the previous behaviors, the behavioral measure in the previous block should not be a median value when the behavioral difference is zero (red fitted line in Figure 2D). This indicates that the subjects shift their behaviors regarding how they performed in the previous block.

### Bayesian reliability estimation

We used a simple empirical Bayesian approach to compute the reliability of each system (Lee et al., 2014). The estimation of the reliability of the method was based on the direct experience of the RL agent, that is, the state-prediction error (SPE) and reward prediction error (RPE) from the MB and MF, respectively. If the average prediction error of a learner (MB or MF) is close to zero, the agent considers the corresponding learning strategy to be more reliable, thereby allocating more behavioral control to it. Specifically, we computed the posterior probability of the system making zero prediction errors (0-PEs) based on the assumption that the parameter of distribution of the 0-PE set was drawn from the Dirichlet prior (Koller and Friedman, 2009). The posterior distribution was subsequently translated onto the reliability value using the mean-variance ratio (Janesick, 2001; Ma et al., 2006; Pennini and Plastino, 2010).

### Online estimation of prediction error distribution

The above-mentioned Bayesian reliability module required an estimation of the 0-PE. Note that the strategy control model reported in the previous study used threshold parameters to classify the prediction error signal into 0-PEs or non 0-PEs (Figure 3B). However, to incorporate dynamic changes of the task structure into 0-PE, we used a DPGMM (Maceachern and Müller, 1998; Neal, 2000; Rasmussen, 2000). The DPGMM is a non-parametric online clustering model that performs updates of the posterior distribution of the incoming data. We set a cluster nearest the zero point as the 0-PE distribution by comparing absolute mean of the PE distribution of every cluster. However, for SPE, there is no need for its absolute value because it ranges between zero and 1. Owing to the online learning capability of the DPGMM, the 0-PE distributions were continually updated to best account for the patterns of the prediction error that the agent currently experiences. Refer to Supplementary Section A for full details.

### Strategy control to minimize lower bound of the prediction error

Finally, to compute the model choice probability *w*, we used a two-state transition mode (Dayan and Abbott, 2001), incorporating the following dynamic changes to the reliability:

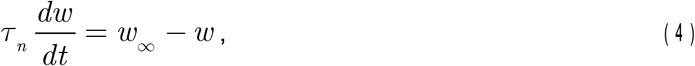

where the time constant (τ) and steady state (*w*_∞_) were computed using the two transition rates (MB→MF, MF→MB); in addition, the reliability was an input in the transition rate function.

It must be noted that the strategy control model in the previous report (Lee et al., 2014) used the above full dynamic transition, whereas the adaptive control model used only a steady-state representation assuming that the dynamic changes in the environment were already incorporated into the reliability:

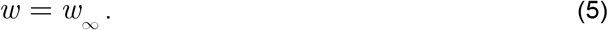

The model integrates the MB and MF learning system by computing the sum of the state-action value Q weighted by the model choice probability, *w* :

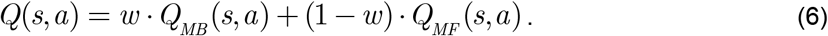

The model uses stochastic action selection with a SoftMax function (Gläscher et al., 2010; Luce, 2012). To evaluate the predictability of the model, we computed the negative log-likelihood of the action selection probability of the observed choices, which were summed over all trials for each participant.

### Model comparison

We tested six different models. Two of them were versions that used only one learning strategy without arbitration (MB and MF). The third version, a strategy control model, is the computational model that previously showed the best performance (Lee et al., 2014). This model achieved a Bayesian reliability estimation and a Pearce–Hall reliability estimation for MB and MF, respectively, and subsequently used the two-state dynamic transition process to integrate reliability information. The last three versions were DPGMM-based models equipped with an online estimation of zero prediction error. These three models used the Bayesian reliability estimation for both MB and MF (a full adaptive control model), for MB only (an adaptive control model for SPE), and for MF only (an adaptive control model for RPE), respectively. We used the Nelder–Mead simplex algorithm (Lagarias et al., 1998) for optimization. We randomly generated seed parameters and subsequently performed the optimization 200 times.

Because DPGMM-based models update the prediction error distributions on each trial, we expected that they could capture changes in the dynamics of the task context. It must be noted that the strategy control model required a dynamic transition to compute the model choice probability, *w*, whereas the full adaptive control model was expected to dispense this part as the model itself already accounted for the dynamics through the online estimation of the 0-PE signals. Refer to Supplementary Table 2 for a full comparison of the free parameters.

There are some technical reasons why the DPGMM-based model demonstrated a much better performance. First, this version is simpler than the strategy control model (smaller number of free parameters) because of its non-parametric clustering capability. We only needed to fit two free parameters in the full adaptive control model. Second, the DPGMM-based model adequately reflects the way individuals perceive their own mistakes (prediction error). The Lee2014 model divided all prediction error data into three different categories (i.e., positive, zero, and negative) using thresholds. However, the DPGMM-based model does not have this constraint. This means that the DPGMM-based arbitration model is more suitable for reflecting the individual effects. Third, the DPGMM-based model adaptively changes the 0-PE distributions. This enables an adaptive reliability estimation of RL learners, based on changes in the average amount of prediction error contingent upon changes in the task context.

### Source of fMRI data

We re-analyzed the fMRI data used in a previous report (Lee et al., 2014). A total of 22 participants were scanned on a 3T Siemens (Erlangen) Trio scanner located at the Caltech Brain Imaging Center (Pasadena), with a 32-channel RF coil for all scanning sessions. Details of fMRI data acquisition can be obtained from the method section of the paper by Lee et al. (2014).

To verify that the computational model reliably recovered the behavioral patterns for strategy control between the MB and MF, we plotted the choice optimality for four combinations of task context, and conducted a GLM analysis with the task context as independent variables and choice optimality as the dependent variable. For the analysis, we actively sampled 40,000 trials for each subject from the computational model. There are four candidate models: MB, MF, Lee2014 (the previous model (Lee et al., 2014)), and our best model. We also compared the effect of the task context on the choice optimality with the behavioral effect, in terms of how much of the proportion was included in the reliable behavioral effect range, which is defined by the mean effect size ±SEM.

### Whole-brain fMRI analysis

The SPM12 software package was used to analyze the fMRI data (Wellcome Department of Imaging Neuroscience, Institute of Neurology, London). The preprocessing protocol was the same as that used in previous research. Briefly, the first four volumes of images were discarded to avoid T1 equilibrium effects, slice-timing of the functional images were corrected to adjust for slight temporal differences within each image, motion-related artifacts were corrected, the image was spatially transformed to match the standard echo-planar imaging template of the brain, and the images were smoothed using a 3D Gaussian kernel (6-mm FWHM) to account for the differential anatomy of the participants.

To demonstrate the neural correlates of MB/MF RL and arbitration, we applied a general linear model (GLM) with eight regressors in total. The design matrix includes the following regressors: (R1) regressors encoding the average BOLD response at two choice states and one outcome state; (R2, R3) parametric regressors encoding PE signals from MB learner and MF learner, respectively; (R4) a parametric regressor encoding the absolute mean value of the zero RPE (i.e. the RPE baseline); (R5) a parametric regressor encoding the variance of the zero SPE; (R6) a parametric regressor encoding the mean value of the zero SPE (i.e. the SPE baseline); (R7) a parametric regressor encoding the normalized maximum reliability of MB and MF learner; (R8) a parametric regressor encoding the chosen minus unchosen action value of the arbitrator (Q); and (R9, R10) parametric regressor encoding action value estimates of MB and MF learner (QMB, QMF), respectively.

All findings reported in the current paper survived a whole-brain correction for multiple comparisons at the cluster level (p < 0.05 corrected), except for the value signals. For testing the value signals, we used a small-volume correction (SVC) in which the center coordinates were determined based on well-grounded works of highly cited studies on value encoding. Details of voxels that survived the statistical tests are outlined in the supplemental information (Supplementary Table 4). We ran a leave-one-subject-out GLM (Esterman et al., 2010), that is, the method ran 22 GLM analyses with one subject left out in each analysis, and each GLM defined the voxel cluster for the subject left out. The GLM analysis was conducted without serial orthogonalization to eliminate the possibility that the order of regressors in the GLM would interfere with the neural results.

### Bayesian model selection on behavioral and fMRI data

To thoroughly test our computational hypothesis about the encoding of the task structure dynamics by updating prediction error baseline against the previous hypotheses, we ran a Bayesian model selection (BMS) (Stephan et al., 2009) on the behavioral and neural data. The BMS analysis helped us identify the best version of an arbitration model in terms of the tradeoff between the model fit and model complexity for the behavioral and neural data. To conduct the BMS analysis on neural data, we defined the region of interest using the SPM toolbox, MarsBaR (Brett et al., 2002) (http://marsbar.sourceforge.net/). We tested one ROI for updates of the SPE baseline. We created our own ROI to compare the strategy control model and the adaptive control model.

### Psychophysiological interaction (PPI) analysis

To elucidate the functional connectivity between regions participating in the MB and MF RLs and the strategy control, we conducted a PPI analysis (Friston et al., 1997). First, we extracted the first eigenvariate of BOLD signals from the 5-mm sphere of several regions, including the posterior putamen, ventromedial prefrontal cortex, lingual/fusiform gyrus, and bilateral vlPFC. The regions were established as playing key roles in the strategy control and for updates of the prediction error baseline of the RL agents. Second, we extracted psychological variables including the SPE baseline, RPE baseline, and the model choice probability *w*. The final step of the PPI analysis followed the same steps for the whole-brain fMRI analysis using the GLM, but we only have an interaction signal (between the psychological term and the physiological activity) as an interested regressor. However, we included the psychological term and physiological activity without interest to avoid any confound from the main effect.

