## Supplementary information for "Prefrontal solution to the bias-variance tradeoff during reinforcement learning"

Dongjae Kim et al., 2020

**Supplementary section**

**Section A. Online estimation of 0-PE signals using Dirichlet process Gaussian mixture model**

To estimate the zero state/reward prediction error we implemented a Dirichlet process Gaussian mixture model (DPGMM) (Maceachern and Müller, 1998; Neal, 2000; Rasmussen, 2000) During each trial the prediction error was used to update existing prediction error clusters. The prediction error signal (x) is represented by the sum of probability of x, which belongs to each existing cluster (Gaussian mixture model):

$$P(x_i) = \sum_{k=1}^{K} P(c_i = k) P(x_i|\theta_k), \qquad (1)$$

where K is the total number of clusters and $c_i$ is the indicator representing each cluster label. $P(x_i|\theta_k)$ is the Gaussian distribution for the cluster k, whereas $\theta_k$ is a parameter of the Gaussian distribution. This GMM can be extended to an infinite Gaussian mixture model (IGMM) that does not require a specified number of clusters. However, we use the concept of the Dirichlet process mixture model to cluster, without prior information about the number of clusters. This method was referred to as the DPGMM across the whole paper.

$$x_i|c_i = k, \qquad x_i \sim \text{Gaussian}(\theta_k)$$

$$\theta_k \sim G \qquad (2)$$

$$G \sim DP(\alpha, G_0)$$

where G is drawn from Dirichlet process with hyperparameter $\alpha$ and base distribution $G_0$, over Gaussian parameters. Here, $G_0$ determines the prior distribution over the Gaussian parameters. As the conjugate prior of the mean of the Gaussian distribution is also Gaussian, and given the Inverse-Wishart distribution for the covariance matrix (or Wishart distribution for the precision matrix), we can formulate joint distribution of parameters of the Gaussian, Inverse-Wishart distribution.

$$\mu_k \sim \mathrm{Gaussian}(\mu_0|\Sigma_k/\kappa_0)$$

$$\Sigma_k \sim \mathrm{Inverse}-\mathrm{Wishart}(\Lambda_0^{-1}, v_0) \quad (3)$$

$$\mu_k, \Sigma_k \sim \mathrm{Gaussian}, \mathrm{Inverse}-\mathrm{Wishart}(\mu_0, \kappa_0, \Lambda_0^{-1}, v_0)$$

This set of hyper-parameters $(\mu_0, \kappa_0, \Lambda_0^{-1}, v_0)$ represent prior belief of the distribution and clustering of data. More specifically, $\mu_0$ is the prior guess about the mean of the cluster k, and $\kappa_0$ is the pseudo-observations which represent the probability of its occurrence. $\Lambda_0^{-1}, v_0$ represent prior belief about the covariance of each cluster. In the current experiment, there is no need to optimize prior belief as we did not have any prior beliefs.

Posterior prediction by means of collapsed Gibbs sampler Even though we modeled the DPGMM, and set the base distribution as a Gaussian, Inverse-Wishart distribution, we still required the posterior prediction as that was our real focus. The posterior distribution can be formulated in the same format as the representation of the Chinese Restaurant Process(CRP) (Pitman, 2006):

$$p(C|X) = \int P(C, \Theta|X)d\Theta$$
$$\propto P(C)\int P(X|C, \Theta)P(\Theta)d\Theta \quad (4)$$

We integrated out $\Theta$ to yield the expression that only depends on $C$, a set of indicators. Markov chain Monte Carlo (MCMC) methods frequently used to predict posterior, and Gibbs sampling is one of the popular methods. However, previous studies (Bush and MacEachern, 1996; Maceachern and Müller, 1998; Neal, 2000; West and Escobar, 1993)noted that the collapsed Gibbs sampling method is more efficient because it has less computational cost demand (Liu, 2008).

The Collapsed Gibbs sampler is the method that integrating out, not only the latent class parameters, but also a set of indicators. This method is more rapid and efficient because it shrinks the state space of the posterior distribution.

Thus, the collapsed Gibbs sampler samples each data point's indicator using the following:

$$\begin{aligned}
&p(c_i = j|C_{\neg i}, X, \alpha) \propto P(X|C)P(C|\alpha) \\
&\propto P(X|c_i = j, C_{\neg i})P(c_i = j|C_{\neg i}, \alpha) \\
&\propto P(x_i|X_{\neg i}, c_i = j, C_{\neg i})P(X_{\neg i}|c_i = j, C_{\neg i})P(c_i = j|C_{\neg i}, \alpha) \\
&= P(x_i|X_{\neg i}, c_i = j, C_{\neg i})P(X_{\neg i}|C_{\neg i})P(c_i = j|C_{\neg i}, \alpha) \\
&\propto P(x_i|X_{\neg i}, c_i = j, C_{\neg i})P(c_i = j|C_{\neg i}, \alpha) \\
&= P(c_i = j|C_{\neg i}, \alpha)P(x_i|X_{c_i=j,\neg i})
\end{aligned} \tag{5}$$

where $X_{c_i=j,\neg i}$ means all the data that belongs to j-th cluster except i-th data point, $x_i$. Then, using the conjugate prior, the distribution of the $P(x_i|X_{c_i=j,\neg i})$ is student-t with hyperparameters $(\mu_0, \kappa_0, \Lambda_0^{-1}, v_0)$. The update rule of those parameters are as follows:

$$\begin{aligned}
&\mu_k = \frac{\kappa_0}{\kappa_0 + N}\mu_0 + \frac{N}{\kappa_0 + N}y \\
&\kappa_n = \kappa_0 + N \\
&v_n = v_0 + N \\
&\Lambda_n = \Lambda_0 + S + \frac{\kappa_0 n}{\kappa_0 + N}(\bar{x} - \mu_0)(\bar{x} - \mu_0)^T
\end{aligned} \tag{6}$$

Thus, by iterating sampling and updates of parameters and hyperparameters, we can yield clusters without sophisticated parameter optimization. For this collapsed Gibbs sampler, we used MATLAB code that was made for spike sorting by Frank Wood (Wood and Black, 2008).

**Section B. Neural correlates of prediction error baseline**

As our computational hypothesis naturally assumed that the prediction error signal functionally overlapped with its baseline, it would be hard to tease out neural correlates of the mean of zero prediction error distribution (i.e. prediction error baseline; Supplementary Figure 6A) from the area that processes prediction error. One viable alternative is to compare neural predictability between the version of the model with prediction error baseline (the full adaptive control model) and the version without the adaptive control (the strategy control model) because two models yielding different behavioral predictions would essentially have different episodes of events. In doing so, we implemented a BMS analysis (Stephan et al., 2009) to formally test whether the neural prediction of the adaptive control model better accounted for neural activities in the brain areas associated with prediction error and behavioral control, in comparison to that by the strategy control model. Our model demonstrated dominant neural activities in the region of fusiform/lingual gyrus, which are the brain areas that have been indicated to encode SPE baseline (Supplementary Figure 6B). However, we could not conduct BMS analysis for RPE baseline because not only RPE baseline but multiple signals (including SPE, the uncertainty of zero SPE, and value computation of MF) had also a significant correlation with the brain area including the pre-SMA.

**Section C. Functional connectivity of strategy control by update of prediction error baseline**

To further understand the functional effects of adaptive control (i.e. updates of prediction error baseline) on strategy control, we conducted psychophysiological interaction (PPI) analysis (Friston et al., 1997). We tested the effect of psychological variables, including SPE baseline, RPE baseline, and the degree of MB control when the brain combines MB and MF systems (i.e. $w$ in $v(s;w) = wv_{\mathrm{MB}}(s) + (1-w)v_{\mathrm{MF}}(s)$ ) (Supplementary Figure 7A), while physiological variables were extracted from bilateral vlPFC, posterior putamen, and vmPFC. As a result, we found four meaningful paths that were modulated by the psychological terms (Supplementary Figure 7B).

First, the correlation between the neural activity pattern of posterior putamen and that of vlPFC decreases with the degree of MB control ($w$) and the PE baseline of the MB system (SPE baseline), and increases with the PE baseline of the MF system (RPE baseline). Specifically, the degree of neural connectivity between posterior putamen and left vlPFC increases with the RPE baseline (cluster-level FWE corrected $p<0.05$), whereas it decreases with the SPE baseline and the degree of model-based control (after small-volume correction [SVC], peak-level FWE corrected $p<0.05$). Also, the degree of neural connectivity between posterior putamen and right vlPFC decreases with the SPE baseline and the degree of MB control (after SVC, peak-level FWE corrected $p<0.05$). Second, the correlation between the neural activity pattern of posterior putamen and that of vmPFC decreases with the RPE baseline, and increases with the SPE baseline and the degree of MB control (after SVC, peak-level FWE corrected $p<0.05$). Third, the correlation between the neural activity pattern of left vlPFC and that of vmPFC decreases with the RPE baseline (after SVC, peak-level FWE corrected $p<0.05$) The functional connection supplements the previous finding (Supplementary Figure 6), by providing evidence demonstrating how the strategy control was affected by PE baseline. For the last, the correlation between the neural activity pattern of LG/FFG and that of posterior putamen increases with SPE baseline (cluster-level FWE corrected $p<0.05$, see

Supplementary Table 5 for voxel-level results of PPI analysis). Since LG/FFG is the region correlated with SPE baseline, this functional connection provides direct evidence of how SPE baseline works on value computation. We demonstrated how SPE baseline and RPE baseline work on the strategy control. Prediction error baseline of MB and MF has an antagonistic relationship in value computation (posterior putamen), while only RPE baseline was significant on value integration.

**Supplementary figures**

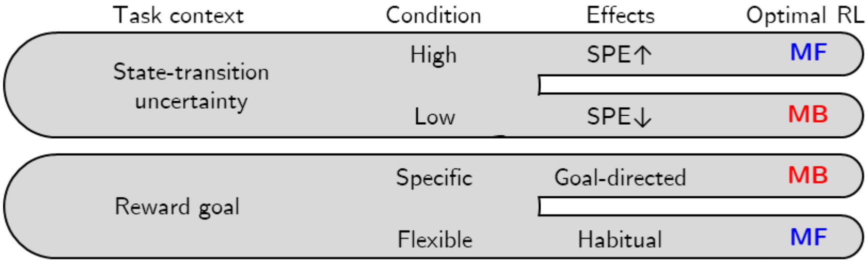

**Figure 1. The optimal RL strategy due to task context**

Each task context is designed to induce one of the RL strategies as the optimal one. Changes in state-transition uncertainty directly modulate the extent of SPE, and changes in reward goal motivate a strategy control by emphasizing the need for precise estimation of state-transitions ("Goal-directed") or simply maximizing reward without it ("Habitual").

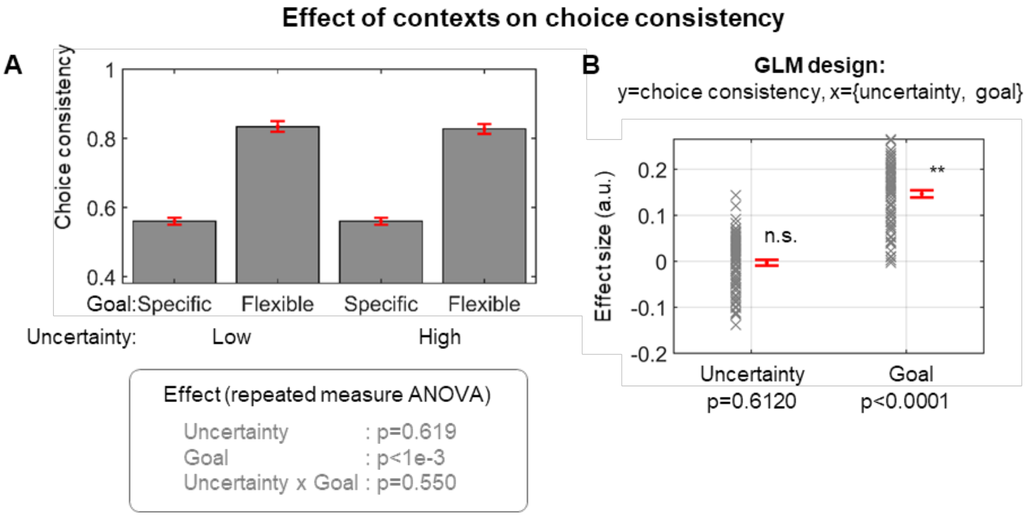

**Figure 2. Behavioral results – choice consistency.**

(A) Choice consistency due to task context. Results of 2-way repeated measures ANOVA are shown in the box (n=82). (B) Results of general linear model analysis. Reward goal has a significant effect on choice consistency (paired t-test; $p<0.0001$), whereas uncertainty does not affect (paired t-test; $p=0.6120$). Error bars are SEM across subjects.

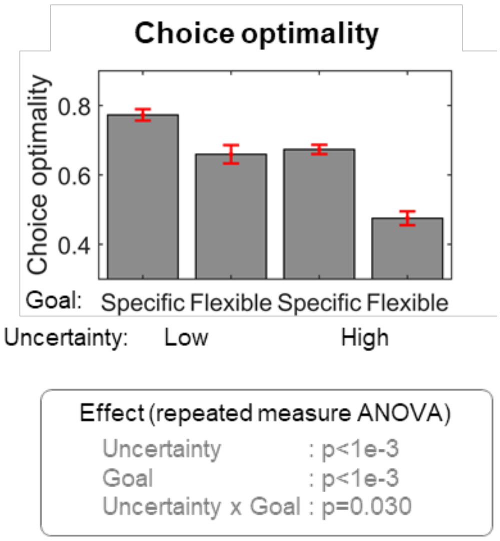

**Figure 3. Behavioral results – choice optimality.**

Choice optimality due to task context. Results of 2-way repeated measures ANOVA are shown in the box (n=82).

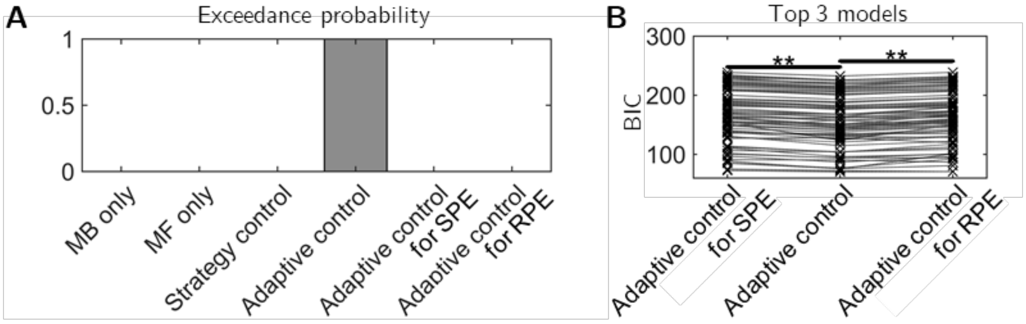

C

| | MB only | MF only | Strategy control | Adaptive control | Adaptive control for SPE | Adaptive control for RPE |
|---|---|---|---|---|---|---|
| # of free parameters | 2 | 2 | 6 | 2* | 3 | 3 |
| AIC | 409.10 | 1345.82 | 157.84 | 151.78* | 155.32 | 157.17 |
| BIC | 415.25 | 1351.96 | 176.39 | 157.89* | 164.59 | 166.32 |

The median values of 82 subjects is shown due to the skewed distribution

**Figure 4. Model comparison**

(A) Bayesian model selection (BMS) result among five different models. (B) Paired-sample t-test result on Bayesian information criterion (BIC) scores (Left: $t_{81} = 15.2470$, $p = 0.000$; Right: $t_{81} = -11.9933$, $p = 0.000$). (C) Details of model comparison (number of free parameters, AIC, BIC scores).

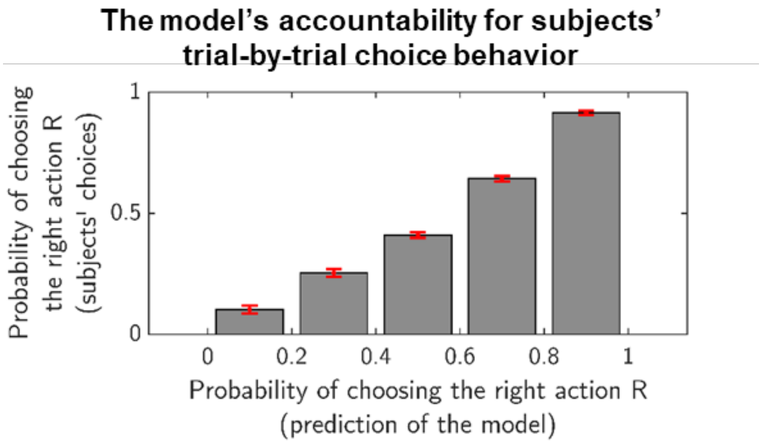

**Figure 5. The proportion of the participants' right action (action R) with the model-predicted probability of choosing the right action.**

The proportion of choosing the action R increased with the probability of choosing the action R predicted by the model, suggesting that the adaptive control model accounted well for choice behavior.

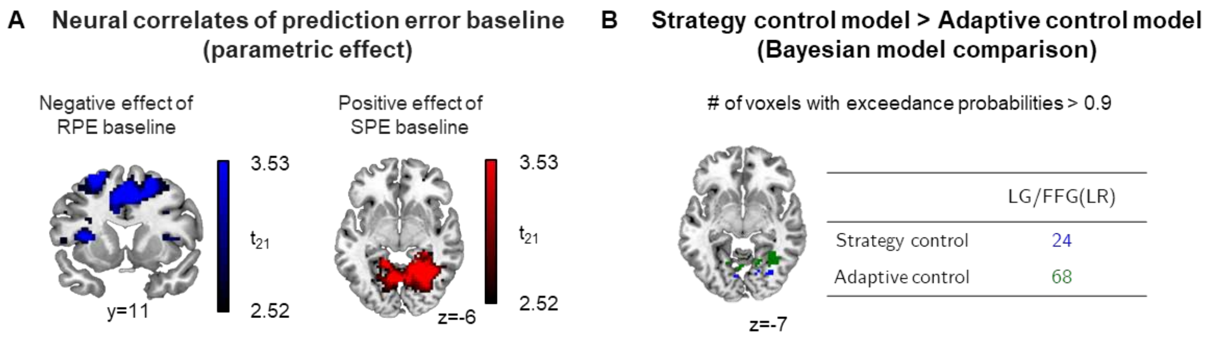

**Figure 6. Neural evidence of prediction error baseline.**

(A) A significant main effect of the RPE baseline was found in the pre-supplementary motor area (pre-SMA) and the significant main effect of SPE baseline was found in the Lingual gyrus (LG) and fusiform gyrus (FFG). See Supplementary Table 4 for full details. (B) Bayesian model selection for the SPE baseline. Bayesian model selection on LG/FFG region of interest showed that the adaptive control model better accounts for the SPE baseline.

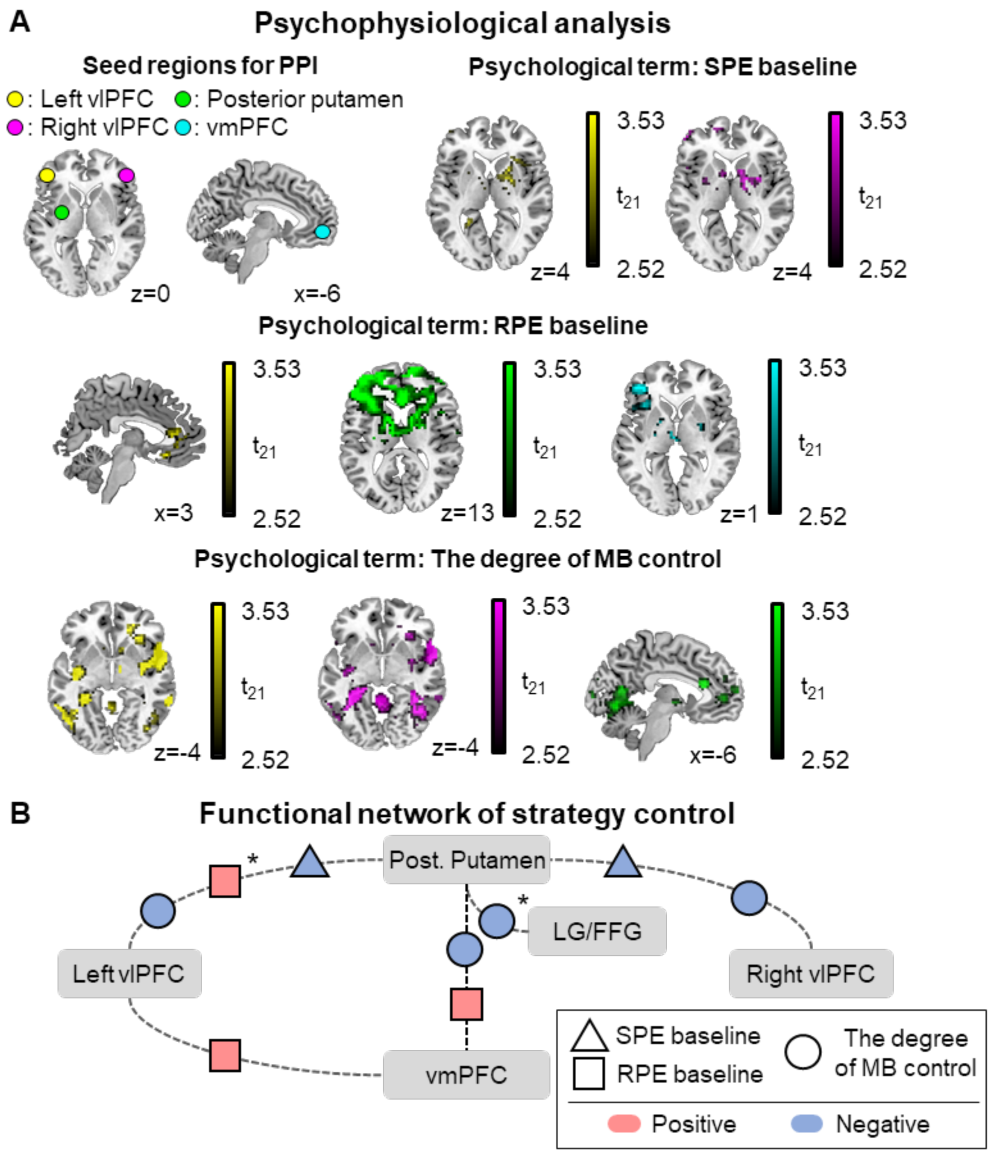

**Figure 7. Functional network analysis.**

(A) Results of PPI analysis. The seed regions include posterior putamen (MF valuation), vmPFC (value integration), and bilateral vlPFC (strategy control). We used the SPE baseline, the RPE baseline, and the degree of MB control as psychological variables. (B) Summary of the PPI analysis. The red and blue color indicates a positive and negative modulation effect, respectively. See Supplementary Table 5 for full details.

### Supplementary tables

#### Table 1. Results of two-way repeated measure ANOVA (behavioral data)

| Source | Uncertainty | Complexity | Type III sum of squares | Degree of freedom | Mean square | F | p |
|---|---|---|---|---|---|---|---|
| **Uncertainty** | Linear | | 1.650 | 1 | 1.650 | 31.803 | .000 |
| **Error (Uncertainty)** | Linear | | 4.202 | 81 | .052 | | |
| **Reward goal** | | Linear | 1.989 | 1 | 1.989 | 96.762 | .000 |
| **Error (Reward goal)** | | Linear | 1.665 | 81 | .021 | | |
| **Uncertainty * Reward goal** | Linear | Linear | .146 | 1 | .146 | 4.901 | .030 |
| **Error (Uncertainty * Reward goal)** | Linear | Linear | 2.413 | 81 | .030 | | |

**Table 2. Free parameters of models**

The first two parameters are required for the reliability estimation of the strategy control model, whereas the third and fourth parameters are required for implementation of the dynamic two-state transition model. The fifth and sixth parameters are SoftMax function parameters for action selection and the learning rate of reinforced learning (RL).

| | ***MB*** | ***MF*** | ***Strategy control*** | ***Adaptive control*** | ***Adaptive control for SPE*** | ***Adaptive control for RPE*** |
|---|---|---|---|---|---|---|
| Number of free parameters | 2 | 2 | 6 | 2 | 3 | 3 |
| Tolerance level of zero SPE | O | X | O | X | X | O |
| Learning rate of absolute RPE | X | O | O | X | O | X |
| MB to MF transition rate | X | X | O | X | X | X |
| MF to MB transition rate | X | X | O | X | X | X |
| Inverse SoftMax temperature | X | X | O | O | O | O |
| Learning rate of the MB and the MF | O | O | O | O | O | O |

MB, model based; MF, model free; RPE, reward prediction error; SPE, state prediction error

**Table 3. Results of two-way repeated measure ANOVA (adaptive control model's data)**

| Source | Uncertainty | Complexity | Type III sum of squares | Degree of freedom | Mean square | F | p |
|---|---|---|---|---|---|---|---|
| **Uncertainty** | Linear | | .068 | 1 | .068 | 6.858 | .011 |
| **Error (Uncertainty)** | Linear | | .798 | 81 | .010 | | |
| **Reward goal** | | Linear | .303 | 1 | .303 | 178.035 | .000 |
| **Error (Reward goal)** | | Linear | .138 | 81 | .002 | | |
| **Uncertainty * Reward goal** | Linear | Linear | .035 | 1 | .035 | 8.040 | .006 |
| **Error (Uncertainty * Reward goal)** | Linear | Linear | .351 | 81 | .004 | | |

**Table 4. Neural correlates**

| X | Y | Z | Hemi. | Peak in the region | P (FWE corrected) | k | Z | T |
|---|---|---|---|---|---|---|---|---|
| **State prediction error (SPE)** | | | | | | | | |
| 36 | 29 | 7 | R | Insula | 0.000* | **295** | 6.30 | 11.12 |
| -30 | 17 | -2 | L | Insula | 0.000* | **180** | 6.30 | 11.09 |
| -36 | 5 | 28 | L | lPFC | 0.001* | **64** | 5.49 | 8.36 |
| -12 | -13 | 4 | L | Thalamus | 0.002* | **20** | 5.27 | 7.75 |
| -33 | -55 | 49 | L | IPL | 0.002* | **71** | 5.26 | 7.73 |
| 12 | 2 | -2 | R | Globus Pallidus | 0.002* | **24** | 5.23 | 7.65 |
| 6 | 23 | 49 | R | SMA | 0.002* | **49** | 5.21 | 7.60 |
| -27 | -73 | 31 | L | IPS | 0.003* | **20** | 5.20 | 7.58 |
| -30 | -82 | 19 | L | IPS | 0.005* | **7** | 5.09 | 7.28 |
| 9 | -10 | 7 | R | Thalamus | 0.008* | **17** | 4.98 | 7.01 |
| 6 | -28 | -2 | R | Thalamus | 0.011* | **8** | 4.91 | 6.84 |
| **Reward prediction error (RPE)** | | | | | | | | |
| 12 | 5 | -11 | R | Ventral striatum | 0.008* | **7** | 4.96 | 6.96 |
| -9 | 5 | -8 | L | Ventral striatum | 0.000+ | 326 | 4.19 | 5.33 |
| -30 | -4 | 1 | L | Dorsal striatum(Putamen) | 0.000+ | 326 | 4.24 | 5.41 |
| -24 | -1 | -5 | L | Amygdala | 0.001+ | 326 | 3.90 | 4.81 |
| 27 | -7 | -5 | R | Amygdala | 0.004+ | 209 | 4.02 | 5.01 |
| 27 | -10 | 10 | R | Dorsal striatum(Putamen) | 0.004+ | 209 | 3.77 | 4.57 |
| **Maximum reliability (*Maximum [Reliability$_{MB}$, Reliability$_{MF}$]*)** | | | | | | | | |
| -51 | 32 | 4 | L | vlPFC | 0.012+ | 112 | 4.61 | 6.17 |
| 48 | 41 | 1 | R | vlPFC | 0.000+ | 217 | 4.33 | 5.60 |
| 12 | 50 | 22 | R | FPC | 0.004+ | 142 | 4.13 | 5.22 |
| **Maximum reliability * SPE baseline** | | | | | | | | |
| 12 | 50 | 22 | R | FPC | 0.036+ | 82 | 4.25 | 5.43 |
| **RPE baseline (negative effect)** | | | | | | | | |
| 0 | 11 | 49 | L/R | Pre-SMA | 0.008* | **24** | 5.07 | 7.23 |
| -24 | 2 | 58 | L | MFG | 0.037* | **2** | 4.68 | 6.32 |
| -54 | -28 | 40 | L | IPL | 0.000+ | 1014 | 4.01 | 5.00 |
| **Uncertainty of zero SPE (negative effect)** | | | | | | | | |
| -54 | -22 | 46 | L | IPL | 0.000* | **58** | 5.60 | 8.68 |
| -6 | 14 | 40 | L/R | dmPFC/SMA | 0.004+ | 174 | 3.98 | 4.94 |
| -27 | -4 | 67 | L | MFG | 0.000+ | 625 | 3.39 | 3.97 |
| **SPE baseline** | | | | | | | | |
| 9 | -52 | -2 | R | LG | 0.019+ | 118 | 3.44 | 4.04 |

| | | | | | | | | |
|---|---|---|---|---|---|---|---|---|
| 36 | -49 | -14 | R | FFG | 0.019+ | 118 | 4.19 | 5.32 |
| **$Q_{MF}$** | | | | | | | | |
| -12 | -4 | 67 | L | SMA | 0.019* | **8** | 4.79 | 6.56 |
| -36 | 38 | 37 | L | dlPFC | 0.004+ | 176 | 4.16 | 5.28 |
| -33 | 56 | 19 | L | alPFC | 0.004+ | 176 | 3.38 | 3.96 |
| -33 | -19 | 7 | L | Posterior putamen | 0.014[1] | - | 3.40 | 3.99 |
| -33 | -1 | -5 | L | Posterior putamen | 0.040[2] | - | 3.01 | 3.41 |
| **$Q_{MB}$** | | | | | | | | |
| 6 | 26 | -17 | L/R | omPFC | 0.028[3] | - | 3.14 | 3.60 |
| **$Q_{ARB}$** | | | | | | | | |
| -6 | 29 | -11 | L/R | vmPFC/OFC | 0.034+ | 88 | 3.88 | 4.77 |
| 0 | -46 | 31 | L/R | PCC | 0.001+ | 219 | 3.87 | 4.75 |

PCC: posterior cingulate cortex; IPS, Intraparietal sulcus; SMA, support motor area; IPL, Inferior parietal lobule; lPFC, lateral prefrontal cortex; MFG, Middle frontal gyrus; FPC, Frontopolar prefrontal cortex; STG, Superior temporal gyrus; om/dl/dm/al/vlPFC: Orbital & medial/Dorsolateral/Dorsomedial/Anterior-lateral/Ventrolateral prefrontal cortex.

[1]: survives after small-volume correction within 10-mm sphere centered coordinates (-27, -13, 4) (Wunderlich et al., 2012) and (-33, -24, 0) (Tricomi et al., 2009)

[2]: survives after small-volume correction within 10-mm sphere centered coordinates (--27, -4, 1) (Lee et al., 2014)

[3]: survives after small-volume correction within 10-mm sphere centered coordinates (0,32,-13) and (3,32,-17) (Wunderlich et al., 2012)

**Table 5. Results of psychophysiological interaction analysis**

| X | Y | Z | Hemi. | Peak in the region | P (FWE corrected) | k | Z | T |
|---|---|---|---|---|---|---|---|---|
| **Negative interaction of right vlPFC by SPE baseline** | | | | | | | | |
| -27 | -1 | 7 | L | Posterior putamen | 0.032[1] | | 3.07 | 3.61 |
| 30 | 2 | 4 | R | Posterior putamen | 0.034[1] | | 3.05 | 3.59 |
| 27 | -4 | 1 | R | Posterior putamen | 0.052[1,2] | | 2.86 | 3.31 |
| **Negative interaction of left vlPFC by SPE baseline** | | | | | | | | |
| 27 | -4 | 1 | R | Posterior putamen | 0.043[1,2] | | 2.95 | 3.44 |
| 24 | -1 | 4 | R | Posterior putamen | 0.044[1] | | 2.94 | 3.42 |
| **Positive interaction of right vlPFC by RPE baseline** | | | | | | | | |
| -6 | 26 | -11 | L | vmPFC | 0.044[3] | | 2.37 | 2.62 |
| **Positive interaction of left vlPFC by RPE baseline** | | | | | | | | |
| 0 | 26 | -14 | L/R | vmPFC | 0.039[4] | | 2.87 | 3.32 |
| **Positive interaction of posterior putamen by RPE baseline** | | | | | | | | |
| -51 | 32 | 7 | L | vlPFC | 0.000+ | 660 | 5.30 | 4.01 |
| -12 | 26 | -2 | L | vmPFC | 0.040[4] | | 2.92 | 3.39 |
| -6 | 29 | -11 | L | vmPFC | 0.046[4] | | 2.85 | 3.29 |
| **Positive interaction of ventromedial prefrontal cortex by RPE baseline** | | | | | | | | |
| -45 | 41 | 4 | L | vlPFC | 0.021[5] | | 3.18 | 3.78 |
| -21 | -1 | -4 | L | Posterior putamen | 0.039[4] | | 2.92 | 3.38 |
| **Negative interaction of right vlPFC by the degree of MB control** | | | | | | | | |
| -33 | -28 | 10 | L | Posterior putamen | 0.028[6] | | 3.16 | 3.63 |
| **Negative interaction of left vlPFC by the degree of MB control** | | | | | | | | |
| 24 | -1 | 64 | R | SMA/MFG | 0.000+ | 687 | 4.40 | 5.74 |
| -33 | -25 | 1 | L | Posterior putamen | 0.047[6] | | 2.94 | 3.32 |
| **Negative interaction of posterior putamen by the degree of MB control** | | | | | | | | |
| -18 | -76 | -2 | L | LG/FFG | 0.003+ | 232 | 3.76 | 4.55 |
| -15 | 23 | -8 | L | vmPFC | 0.024[4] | | 3.15 | 3.61 |

1: survives after small-volume correction within a 10-mm sphere centered coordinate (-27, -4, 1) or (27, -4, 1) (Lee et al., 2014)

2: survives after small-volume correction within a 10-mm sphere centered coordinate (27, -13, 4) (Wunderlich et al., 2012)

3: survives after small-volume correction within a 5-mm sphere centered coordinate (-9, 29, -11) (Lee et al., 2014)

4: survives after small-volume correction within a 10-mm sphere centered coordinate (-9, 29, -11) (Lee et al., 2014)

5: survives after small-volume correction within a 10-mm sphere centered coordinate (-54, 38, 3) (Lee et al., 2014)

6: survives after small-volume correction within a 10-mm sphere centered coordinate (-33, -24, 0) (Tricomi et al., 2009)
